# A five-dimensional functional state space for fingerprinting disease transcriptomes

**DOI:** 10.64898/2026.07.04.736469

**Authors:** Fengyun Nie, Yan Zhuang, Ke Chen, Jiangli Lin, Jianjun Sun

## Abstract

High-throughput transcriptomics has transformed disease biology, but its outputs often remain fragmented into gene and pathway lists that are difficult to compare across conditions or use for human-AI interpretation. We developed a five-dimensional (5-D) functional state space that represents disease transcriptomes as coordinated activity patterns across major biological systems. The framework maps transcriptomic signals onto five functional systems, 14 subcategories, and a distinct infrastructure layer, and was implemented as a reproducible pipeline for functional scoring, cross-condition profiling, benchmarking, and large language model (LLM)-assisted interpretation. Applied to wound healing, sepsis, colorectal cancer-related datasets, an extended GEO atlas of 38 complete case-control disease fingerprints spanning diverse disease contexts, and a TCGA-COAD/READ stage benchmark, the approach recovered interpretable disease-state patterns and retained progression-related information under strong compression. It also improved the quantitative grounding of LLM-generated summaries. This framework provides a compact and auditable representation for comparing disease transcriptomes and supporting human-AI biological interpretation.

**Teaser:** A five-dimensional functional framework turns complex disease transcriptomes into compact, interpretable fingerprints for human-AI analysis.

## Introduction

High-throughput transcriptomic profiling is widely used to characterize disease-associated molecular alterations, identify subtypes, explore pathways, and prioritize biomarkers (*1–5*). However, transcriptomic outputs are often difficult to interpret at a systems level because they are high-dimensional, context-dependent, and fragmented into long lists of genes, pathways, and functional terms (*4*, *6–8*). Organ-level descriptions are clinically intuitive but too coarse to capture cross-system coordination, whereas gene- and pathway-level analyses provide mechanistic detail but are often too granular for compact high-level interpretation. These limitations point to the need for a structured functional state space in which disease transcriptomes can be represented as comparable, interpretable biological states.

The rise of LLMs has created new opportunities for translating structured biomedical outputs into human-readable interpretations (*9–13*). Yet direct use of long gene lists, pathway lists, or enrichment terms can produce context overload, weak quantitative grounding, and unsupported biological generalization (*14–17*). These limitations are especially important for human-AI interpretation, where the input representation strongly influences whether generated summaries remain evidence-based and biologically coherent. A useful intermediate representation should therefore be compact enough for cross-condition comparison and LLM-assisted interpretation, while retaining biologically meaningful structure and traceable quantitative anchors.

This need is particularly relevant to host-response biology, where disease outcomes depend not only on immune-defense intensity but also on repair, resolution, restoration of tissue integrity, and control of collateral injury. Prior host-response modeling by Sun, including the “promoting self-healing and balancing immunity” model and the host-pathogen interaction (HPI) equation, emphasized the dynamic balance among self-healing capacity, immune activity, pathogen/damage burden, and collateral injury (*18*, *19*). This perspective motivated the explicit separation of self-healing/reconstruction and immune defense in the present framework, allowing disease transcriptomes to be fingerprinted not only by immune activation but also by coordinated changes in repair, metabolism, regulation, and continuity-related functions.

In this study, we propose a 5-D functional state space for fingerprinting disease transcriptomes. The framework maps transcriptomic signals onto five major functional systems, 14 functional subcategories, and a System 0 infrastructure layer, and is implemented in a reproducible pipeline linking GEO preprocessing, probe-to-gene mapping, ssGSEA scoring, functional profiling, controlled benchmarking, and LLM-assisted interpretation (*20–23*). We evaluate the framework through internal coherence analyses, wound-healing time-series profiling, sepsis and colorectal cancer case studies, an extended GEO disease fingerprint atlas, a TCGA colorectal cancer stage benchmark, and an LLM input-mode comparison. Together, these analyses test whether disease transcriptomes can be transformed into compact, auditable functional fingerprints that support cross-disease comparison and human-AI biological interpretation.

## Results

### Internal coherence analyses support a non-random functional architecture

The analytical workflow first converts heterogeneous transcriptomic datasets into gene-level matrices and then projects them into the predefined 5-D functional space using ssGSEA-based scoring. The framework defines five parent systems, 14 subcategories, and a System 0 infrastructure layer that was excluded from downstream disease-state functional scoring (**Figure 1**, **Table 1**).

**Figure 1.**
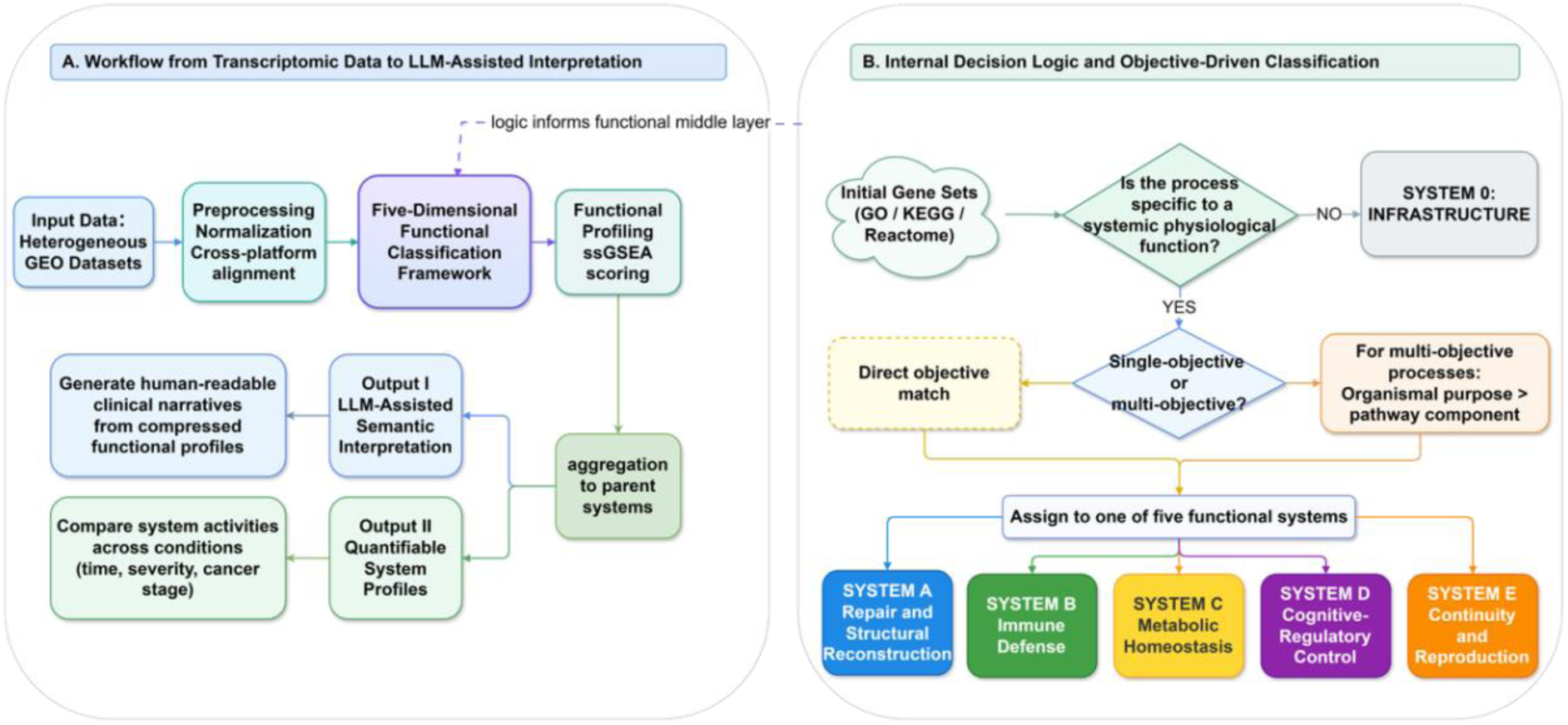
Conceptual and analytical framework of the 5-D functional state-space approach. (A) Overview of the analytical workflow. Heterogeneous transcriptomic datasets are processed through dataset-level preprocessing, gene-level standardization, and probe-to-gene mapping when required. Gene-level expression matrices are then mapped into the predefined 5-D functional framework and quantified using ssGSEA-based scoring. Subcategory scores are aggregated into parent systems to generate functional profiles for cross-condition comparison and LLM-assisted interpretation. (B) Decision logic for objective-driven functional classification. Gene sets are first evaluated as general infrastructure or function-specific programs. Infrastructure processes are assigned to System 0, whereas function-specific processes are assigned to Systems A-E according to dominant physiological objective.

**Table 1.**
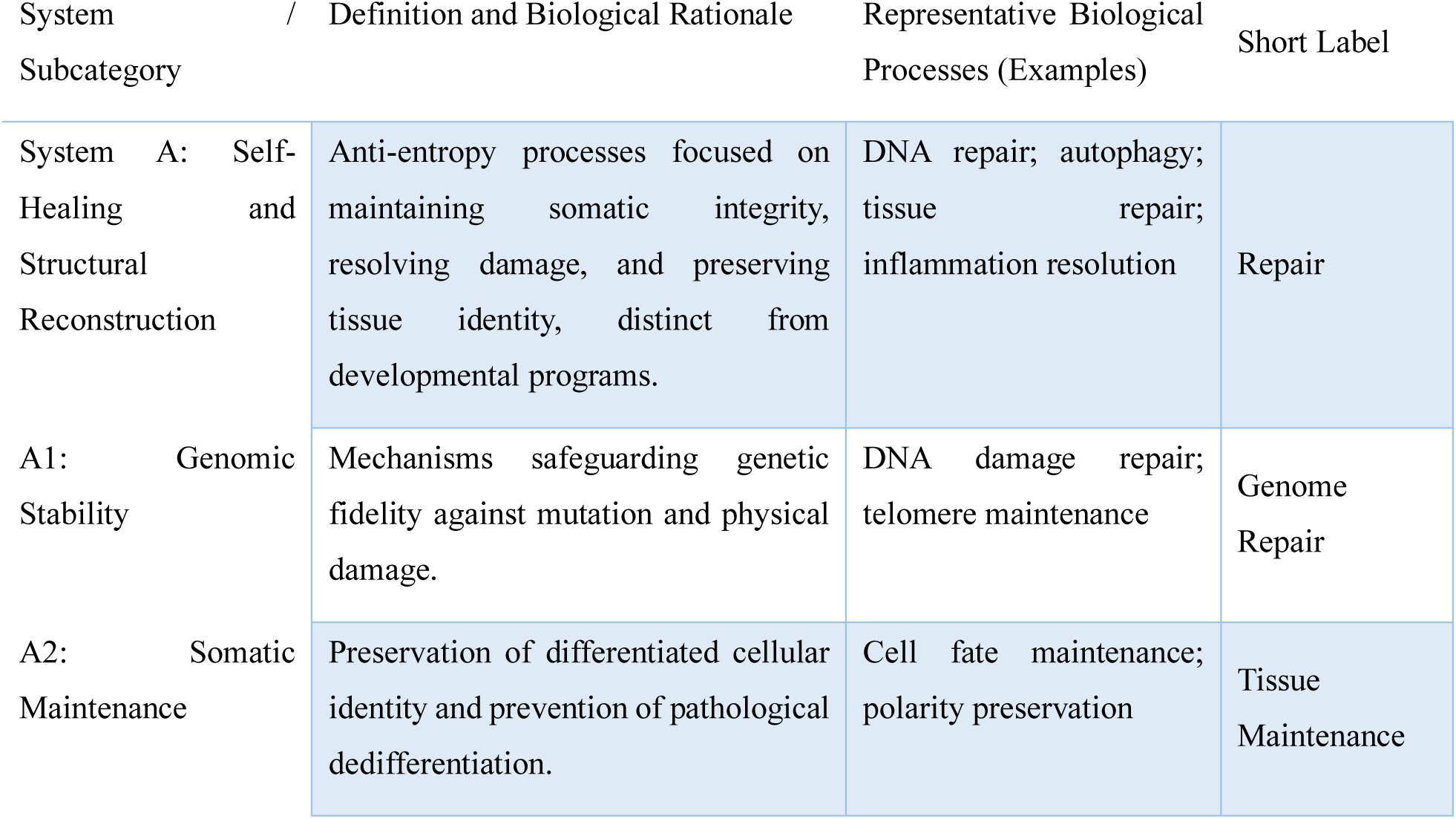
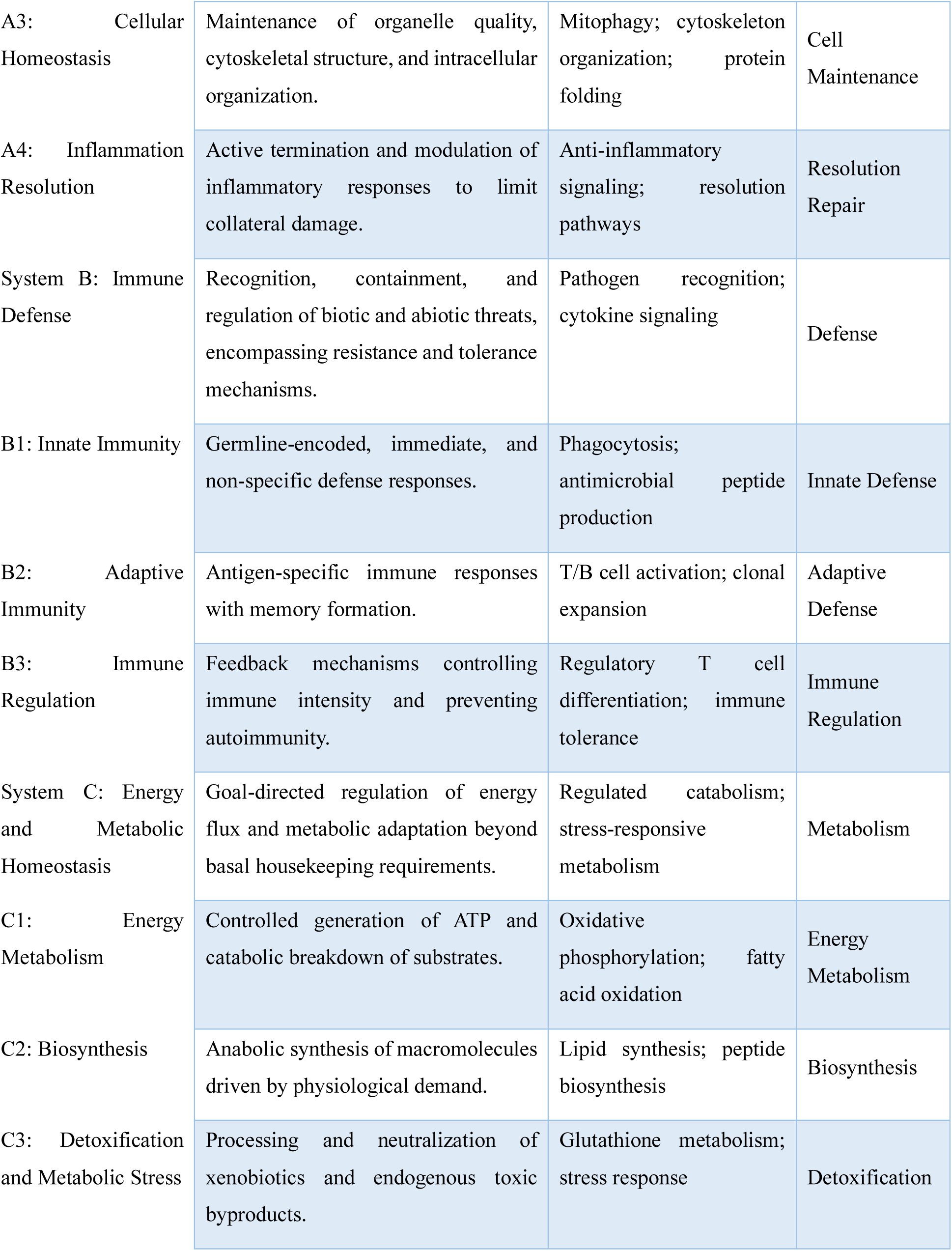
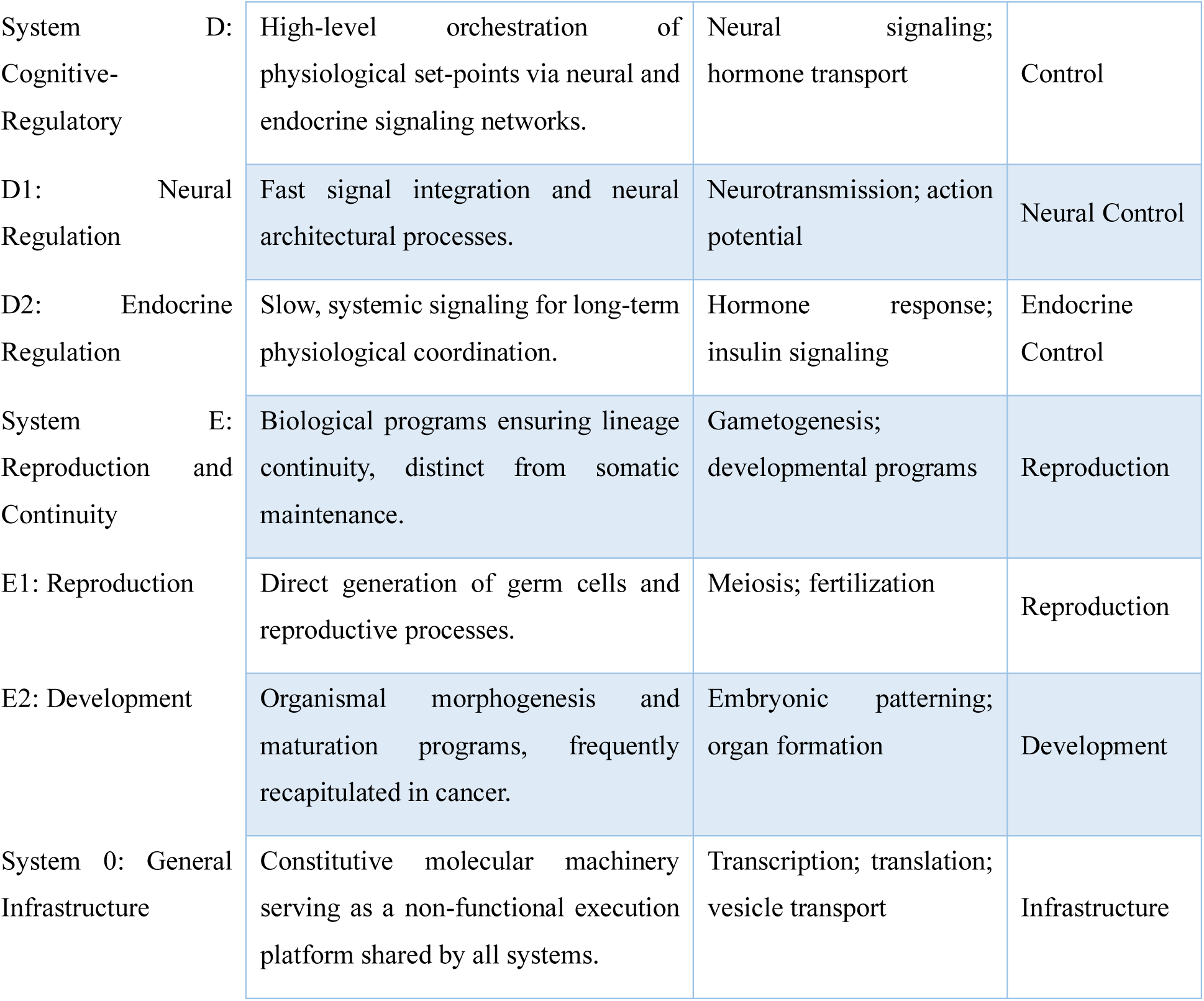
Functional systems and subcategory definitions. This table defines the five parent systems (A-E), 14 functional subcategories, and System 0 infrastructure layer used throughout the study. Systems and subcategories were defined a priori based on dominant physiological objectives rather than inferred from dataset-specific expression patterns. Representative biological processes are illustrative examples rather than exhaustive lists. System 0 denotes constitutive cellular machinery and was excluded from downstream disease-state functional scoring.

Applying the deterministic rules to all GO biological processes and KEGG pathways produced a clear division between System 0 and the five functional systems. Approximately half of all classified biological entities (50.7%) were assigned to System 0, indicating that a large fraction of standard ontology content represents constitutive molecular infrastructure rather than context-dependent physiological objectives (**Figure 2**).

**Figure 2.**
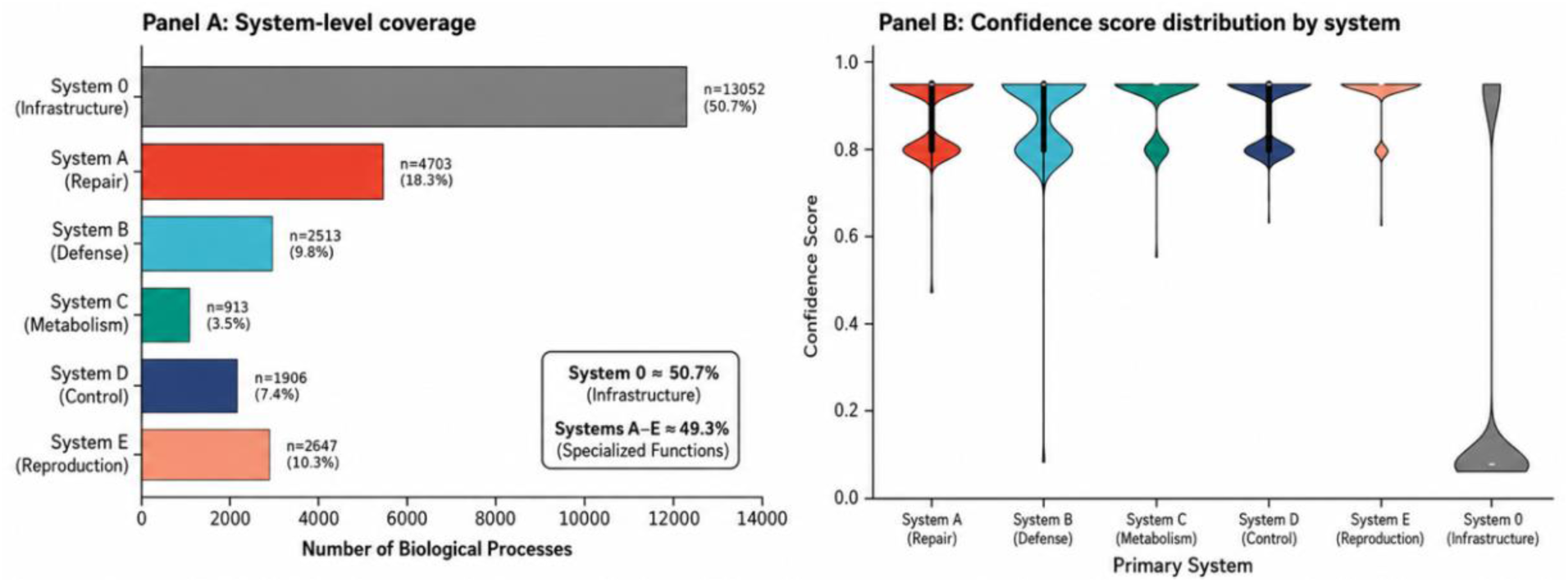
Global distribution of classified GO biological processes and KEGG pathways across System 0 and Systems A-E. **(A)** System-level coverage of classified biological processes across the general infrastructure layer and the five specialized functional systems. Biological processes were assigned using deterministic dominant-objective classification rules. System 0 represents constitutive molecular infrastructure, whereas Systems A-E represent specialized physiological objectives used for downstream disease-state functional profiling. **(B)** Distribution of classification confidence scores for biological processes assigned to each primary system. Violin plots show the score distribution within Systems A-E and System 0. The specialized functional systems showed predominantly high confidence scores, supporting stable assignment of most biological processes to their primary functional objectives.

The five functional systems also showed distinct semantic fingerprints across representative semantic domains. System A was enriched for repair- and structure-related semantics, System B for immune-defense terms, System C for energy and metabolism, System D for signaling and regulatory control, and System E for development and reproduction-related semantics (**Figure 3**). Permutation-based anchor purity and GO graph-proximity analyses further showed that within-system semantic concentration and topological proximity exceeded random expectation, supporting the internal coherence of the objective-driven classification (**Supplementary Figs. S1-S2**).

**Figure 3.**
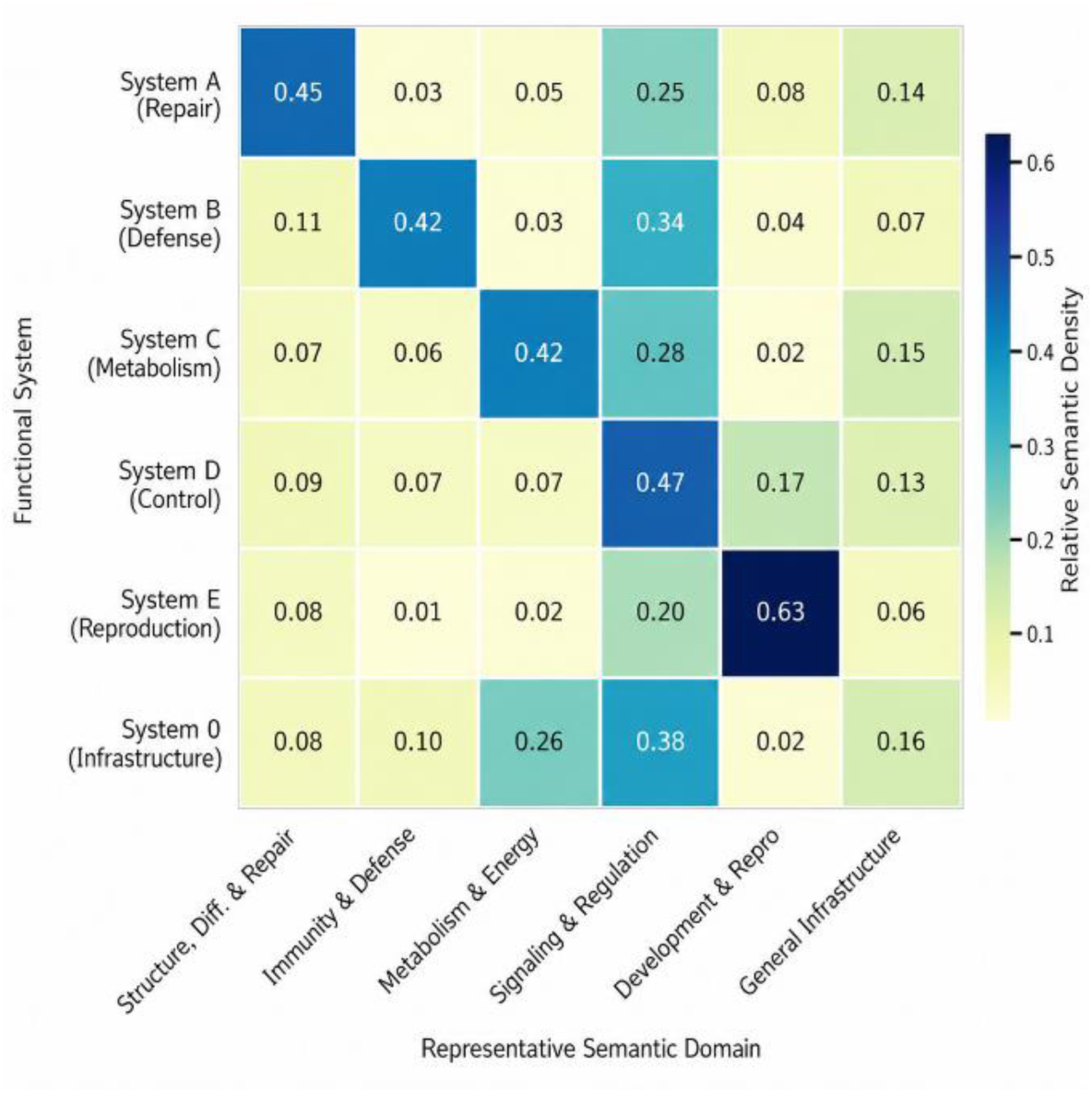
Semantic fingerprints of the functional systems across representative semantic domains. The heatmap shows relative semantic density of each functional system across representative semantic domains derived from anchor terms. Rows correspond to systems, and columns correspond to semantic domains. The panel displays distinct semantic profiles for Repair, Defense, Metabolism, Control, and Reproduction/Continuity systems.

### Wound healing shows staged system- and subcategory-level temporal dynamics

To assess dynamic tissue repair, we analyzed wound-healing time points from baseline to day 21 using the unwounded state as reference. Early time points (baseline to day 7) were derived from GSE28914, whereas later time points (day 14 to day 21) were derived from GSE50425. Because the two windows came from independent cohorts, values were interpreted as relative within-cohort functional changes rather than directly batch-corrected sample-level measurements.

At the system level, all five systems showed a day-1 activation peak followed by attenuation by day 3. Systems D and E showed the strongest day-1 activation, Systems B and C were also induced, and System A showed the lowest day-1 activation. From day 3 to day 7, Systems A, C, D, and E showed secondary increases, whereas System B continued to decline slightly. By days 14 and 21, all five systems had largely returned toward baseline (**Figure 4A**).

**Figure 4.**
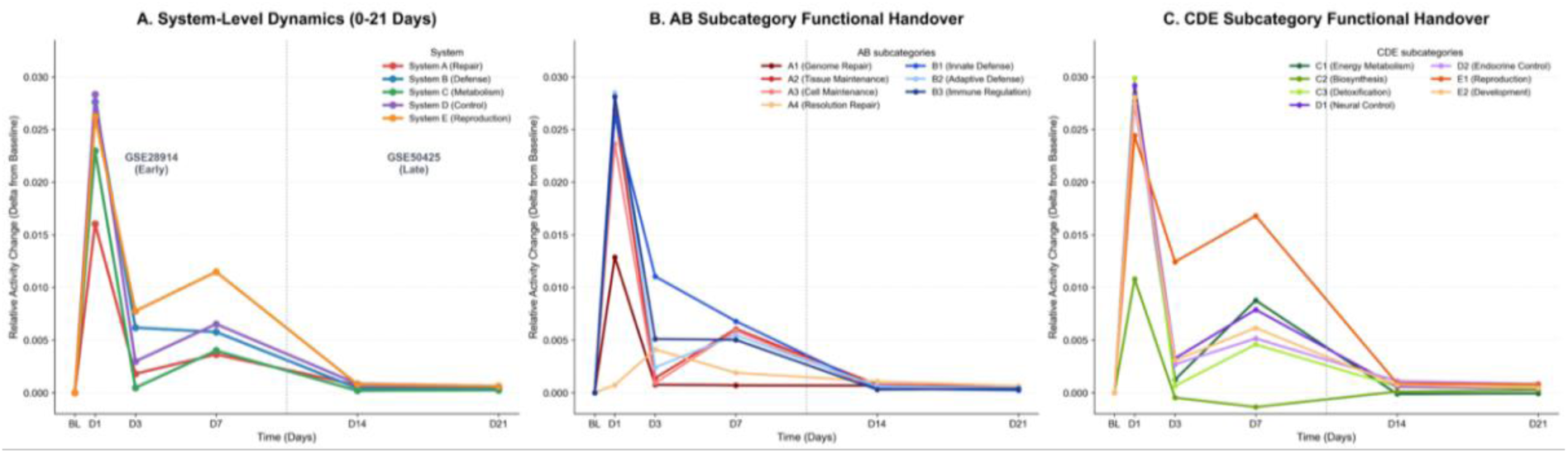
Integrated functional timeline of wound healing across systems and subcategories. **(A)** System-level relative activity changes from baseline to day 21. Early time points (baseline to day 7) were derived from GSE28914, and later time points (day 14 to day 21) were derived from GSE50425; values should be interpreted as relative within-cohort functional changes. **(B)** Subcategory-level trajectories for Systems A and B. **(C)** Subcategory-level trajectories for Systems C, D, and E. Panels display temporal changes in system- and subcategory-level activity during the wound-healing continuum.

Within Systems A and B, B1, B2, B3, A3, and A2 were strongly activated at day 1, whereas A1 showed a more moderate increase. Most A/B subcategories decreased by day 3; A2, A3, B2, and B3 increased again around day 7. A4 showed a delayed pattern, reaching its highest level around day 3 and remaining above baseline through day 7 (**Figure 4B**).

Within Systems C, D, and E, C3, D1, D2, E1, and E2 were strongly induced at day 1, and C2 also increased. Several C/D/E subcategories showed secondary activity around day 7, most prominently E1, with additional delayed increases in C1 and D1. C2 declined rapidly after day 1 and remained near or below baseline thereafter. Phase-specific top-gene analysis showed substantial turnover among the top activated genes contributing to each system, indicating that the system-level trajectories were carried by different gene modules at different phases rather than by one fixed gene set (**Figure 4B, 4C; Supplementary Table S1**). These observations suggest that the 5-D framework can capture dynamic functional handover during tissue repair while retaining gene-level phase specificity.

### Sepsis displays asymmetric metabolic-regulatory activation with reduced blood-based immune-defense signatures

We next analyzed an independent sepsis cohort (GSE65682). Subcategory-level effect sizes showed an asymmetric functional profile rather than uniform shifts across all subcategories. The largest positive shifts occurred in C3 and D2, followed by A3, C1, C2, D1, A4, and E1. In contrast, B1, B2, and B3 showed negative shifts relative to controls, indicating that the strongest transcriptomic differences in this cohort were concentrated in metabolic-stress and regulatory dimensions rather than canonical immune-defense activation (**Figure 5A**).

**Figure 5.**
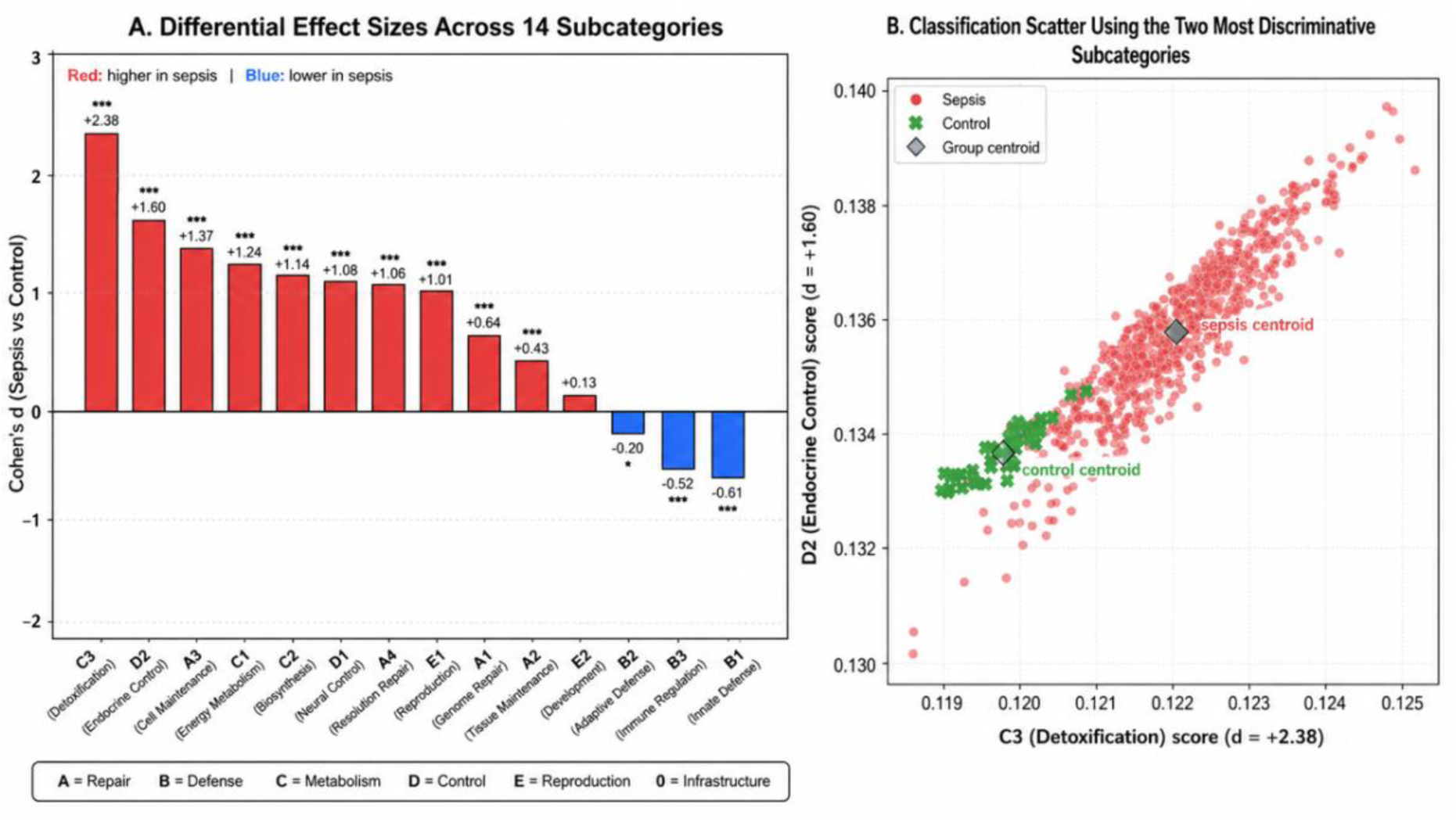
Sepsis-associated subcategory shifts and C3-D2 projection. **(A)** Subcategory-level effect sizes (Cohen’s d) comparing sepsis and control samples in GSE65682. Positive values indicate higher activity in sepsis; negative values indicate lower activity relative to controls. **(B)** Two-dimensional projection of individual samples onto C3 (Detoxification) and D2 (Endocrine Control), the two most discriminative subcategories in this cohort. Because GSE65682 is based on whole-blood leukocyte transcriptomes, immune-defense shifts should be interpreted as blood-based signatures that may reflect both transcriptional state and cell-composition differences.

Projection of individual samples onto C3 and D2 separated sepsis from control samples, with distinct group centroids in this two-dimensional functional space. Thus, metabolic-stress handling and endocrine-regulatory activity contributed prominently to the transcriptomic distinction between sepsis and controls (**Figure 5B**).

Because GSE65682 is based on whole-blood leukocyte transcriptomes, the B1-B3 decrease was treated as a blood-based immune-defense signature that may reflect both leukocyte transcriptional state and cell-composition differences. Within this dataset, however, the coordinated negative shifts of B1-B3 indicate reduced canonical immune-defense signatures in sepsis relative to healthy controls. This pattern suggests dysregulated host-response organization rather than uniform immune activation.

### Colorectal cancer-related datasets reveal recurrent and stage-associated functional patterns

To evaluate robustness across heterogeneous colorectal cancer-related datasets, we analyzed 21 disease-versus-normal comparisons spanning adenoma, carcinoma/tumor, and metastasis contexts. These included 7 adenoma comparisons from 7 datasets (214 disease and 239 control samples), 10 carcinoma/tumor comparisons from 10 datasets (499 disease and 249 control samples), and 4 metastasis comparisons from 4 datasets (153 disease and 110 control samples). Most comparisons used tissue biopsy samples, whereas the carcinoma/tumor group also included peripheral-blood data (**Supplementary Tables S2 and S3**).

Across adenoma, carcinoma/tumor, and metastasis comparisons, A1 and C2 showed the most consistent positive shifts. A4 was also positively shifted across the colorectal cancer spectrum, whereas D2 and C3 showed recurrent negative shifts. These shared patterns were observed despite differences among datasets, platforms, and disease contexts (**Figure 6A-C**).

**Figure 6.**
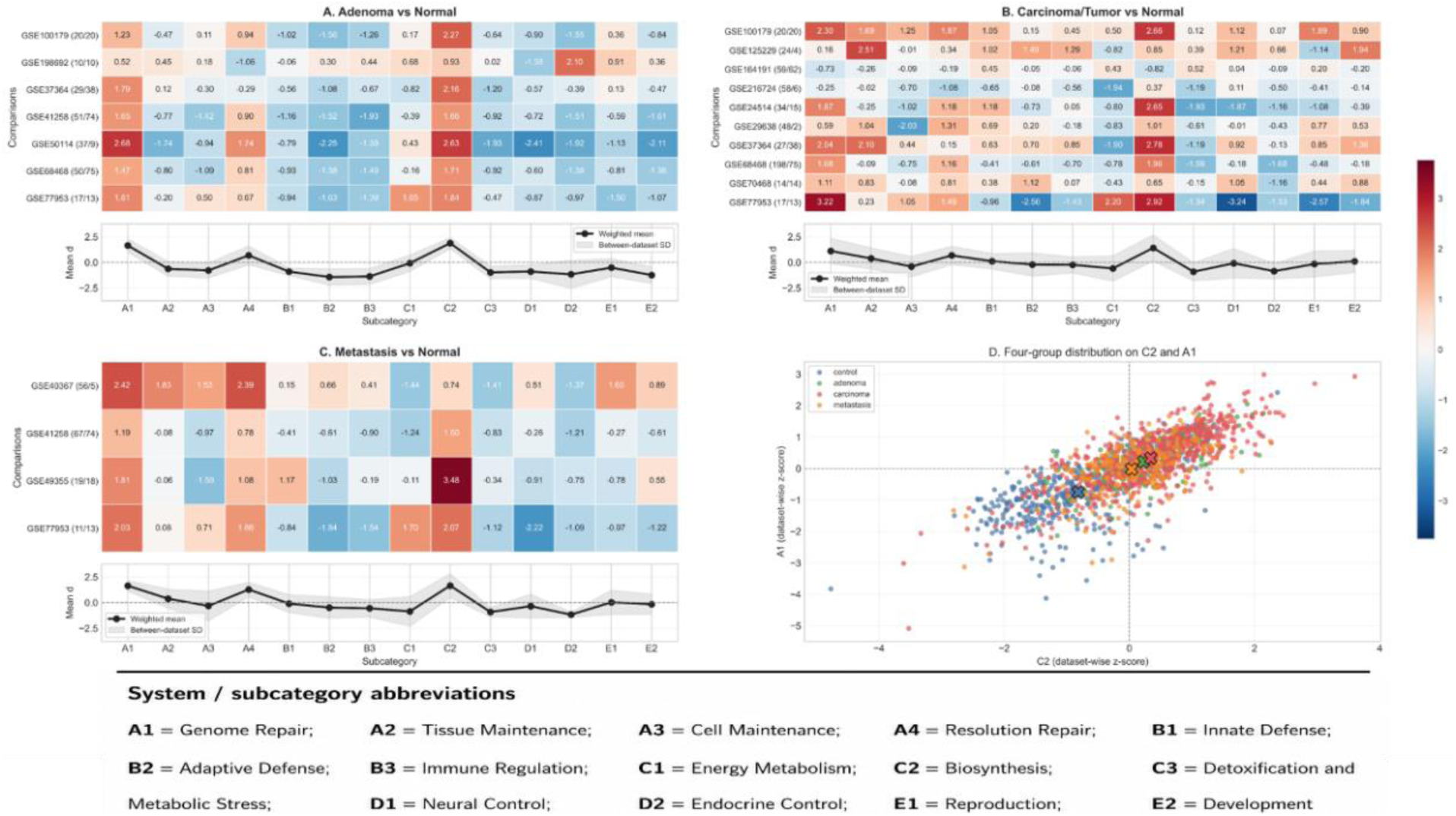
Recurrent subcategory-level patterns across colorectal cancer-related comparisons. **(A-C)** Heatmaps show subcategory-level effect sizes (Cohen’s d, disease versus normal) across independent adenoma, carcinoma/tumor, and metastasis comparisons, respectively. Summary panels show weighted mean effect size and between-dataset variability for each subcategory. **(D)** Sample-level projection onto C2 (Biosynthesis) and A1 (Genome Repair), two recurrent discriminative subcategories across the colorectal cancer-related comparisons.

Stage-associated differences were also observed. Adenoma comparisons already showed elevation of A1 and C2. Carcinoma/tumor comparisons preserved the A1/C2 pattern and showed more evident positive shifts in A2 and A4. Metastasis comparisons retained prominent A4 elevation, while D2 and C3 remained negatively shifted. Immune-related subcategories were more heterogeneous, although B2 and B3 tended to show more reproducible negative shifts than B1. Sample-level projection onto C2 and A1 showed a graded distribution, with controls concentrated at lower values and adenoma-, carcinoma-, and metastasis-enriched samples shifted toward higher values with substantial overlap (**Figure 6D**).

Together, these results show that the 14-subcategory representation captured both recurrent and stage-associated functional patterns across independent colorectal cancer-related datasets. The most consistent shared pattern was A1/C2 elevation, accompanied by A4 positivity and recurrent D2/C3 suppression, while disease-stage groups differed mainly in the relative strengthening of remodeling- and maintenance-associated subcategories.

### The extended GEO atlas supports scalable cross-dataset functional fingerprintin**g**

To evaluate scalability beyond the main case studies, we applied the pipeline to an extended supplementary GEO resource comprising 93 dataset-level analyses across cancer, infection, autoimmune, respiratory, neurodegenerative, metabolic, liver-related, and other disease contexts. For cross-dataset comparison, we focused on 43 realized case-control analyses and defined each disease fingerprint as a 14-dimensional vector of case-versus-control Cohen’s d values (**Figure 7**).

**Figure 7.**
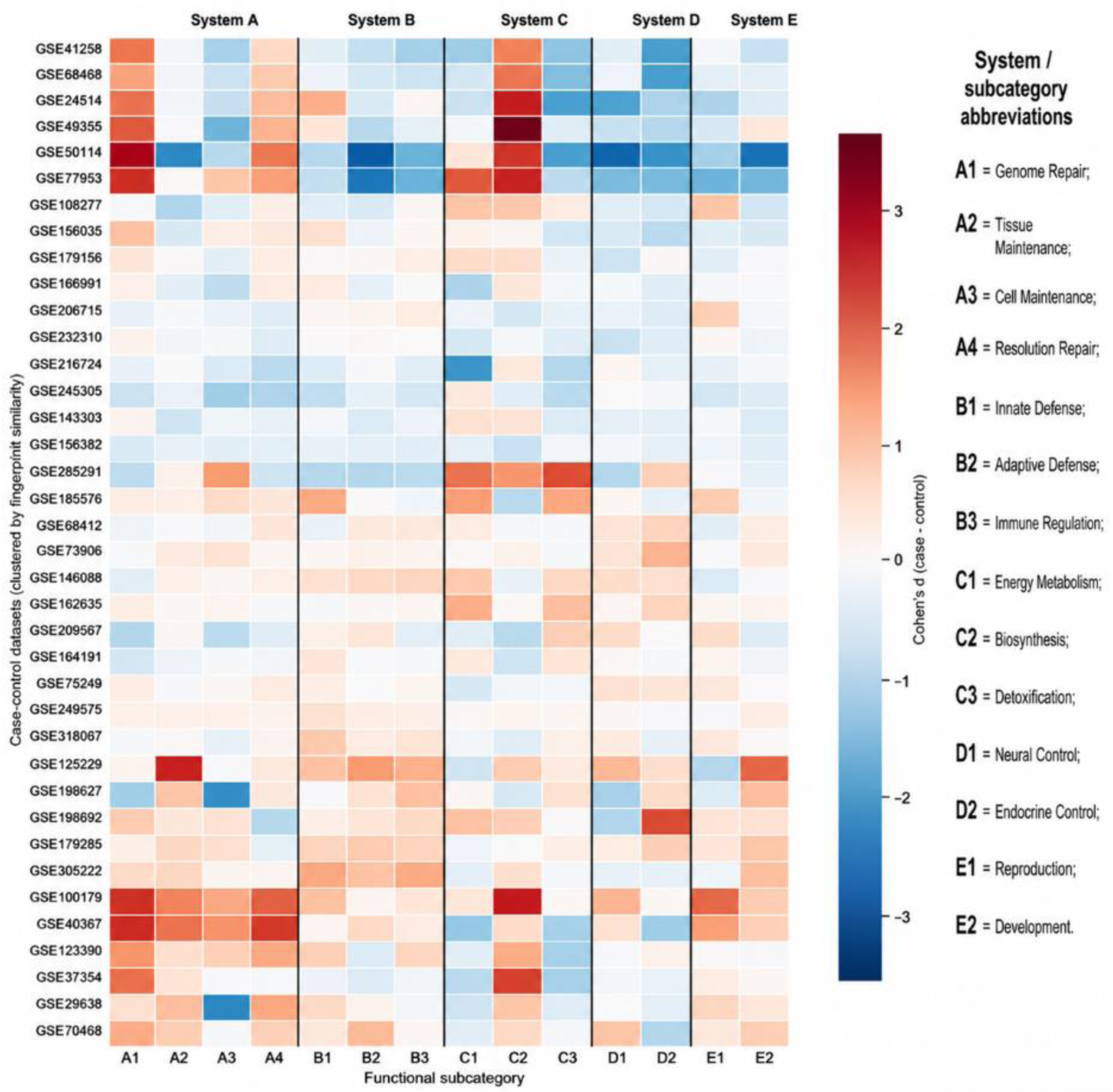
Cross-dataset atlas of 14-subcategory disease fingerprints derived from case-control GEO transcriptomic analyses. Each row represents one case-control dataset, and each column represents one of the 14 functional subcategories. Cell values correspond to case-versus-control Cohen’s d effect sizes, with positive values indicating relative activation in disease and negative values indicating relative suppression. Only datasets with complete 14-subcategory effect-size vectors were included (n = 38).

Among the 43 case-control analyses, 38 contained complete 14-subcategory effect-size vectors and were included in the atlas; five datasets were excluded because one subcategory was unavailable. The included datasets covered 22 cancer datasets, 4 infection datasets, 4 autoimmune datasets, 2 respiratory datasets, 2 neurodegenerative datasets, 1 metabolic dataset, 1 liver-related dataset, and 2 additional datasets. The heatmap showed heterogeneous but comparable disease fingerprints across the 14-subcategory functional space, with both disease-specific patterns and partial recurrence of shared motifs across related conditions (**Figure 7**). These observations suggest that the 14-subcategory representation is sufficiently expressive for scalable cross-dataset functional fingerprinting while maintaining a unified comparative language.

### The 14-subcategory representation retains progression-related information under dimensional compression

To evaluate whether the framework preserved disease-relevant information in a controlled within-disease setting, we analyzed 358 TCGA-COAD/READ samples, including 190 early-stage tumors (stage I/II) and 168 advanced-stage tumors (stage III/IV). The 14-subcategory representation was compared with the five-system representation, Hallmark 50, KEGG pathways, Reactome pathways, GO-slim biological processes, and random 14-gene-set controls using 5-fold cross-validation (**Table 2**).

**Table 2.**
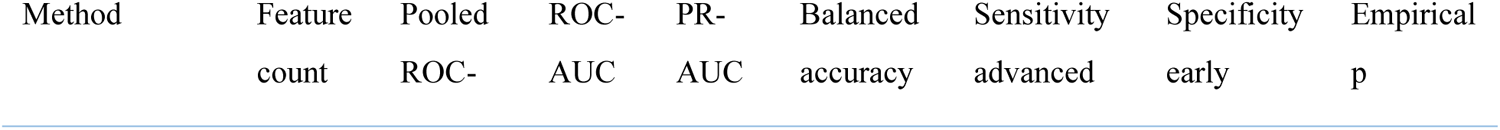
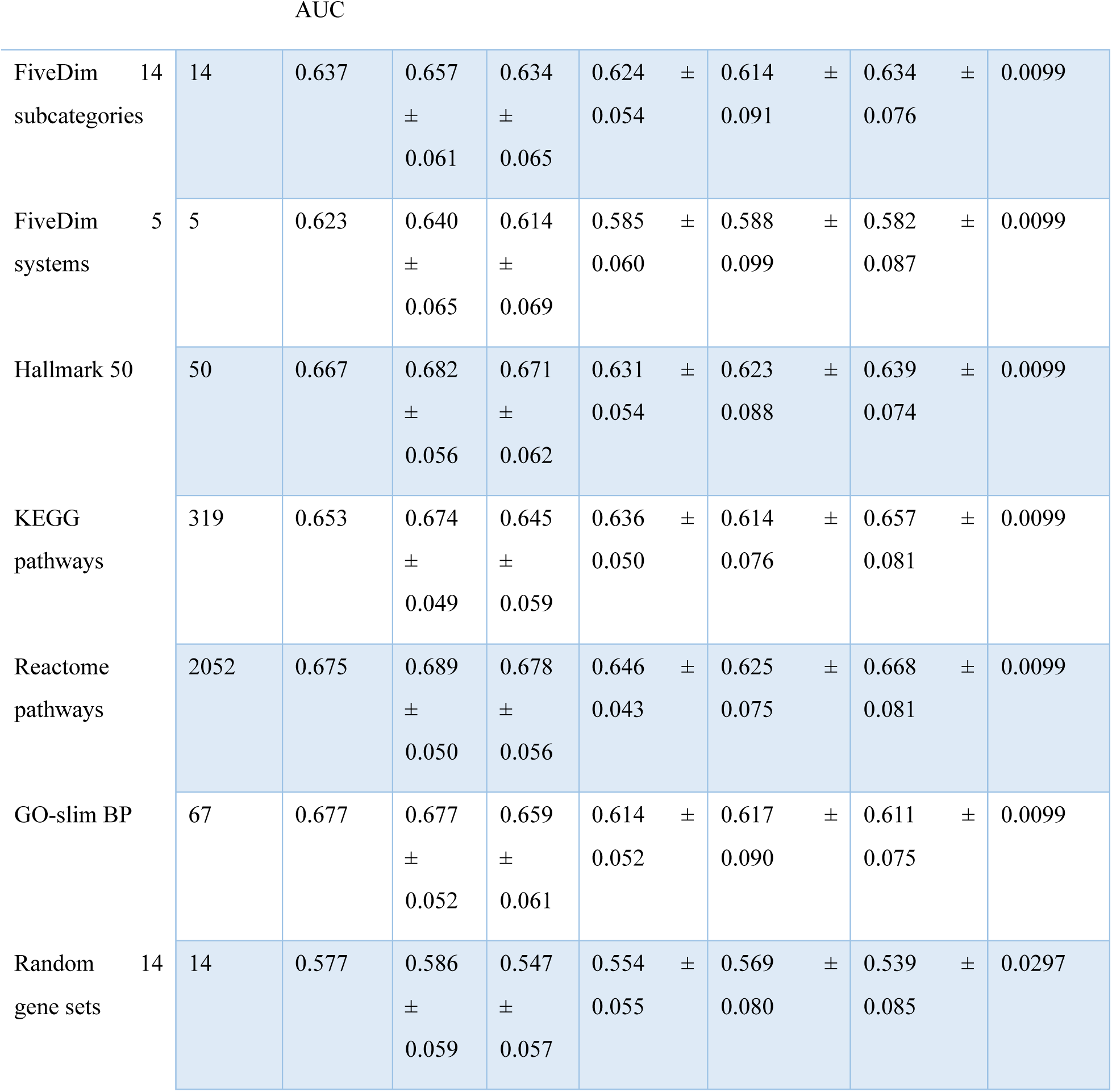
Controlled TCGA colorectal cancer stage benchmark comparing early-stage and advanced-stage tumors. Early-stage tumors were defined as stage I/II, and advanced-stage tumors were defined as stage III/IV. A total of 358 TCGA-COAD/READ samples were included, comprising 190 early-stage and 168 advanced-stage tumors. Each representation was evaluated using 5-fold cross-validation. Pooled ROC-AUC was calculated from pooled cross-validated predictions. Fold-level ROC-AUC, PR-AUC, balanced accuracy, sensitivity for advanced-stage tumors, and specificity for early-stage tumors are reported as mean +/- SD across folds. Empirical p values were estimated by label-permutation testing. The benchmark was designed to evaluate compactness and progression-related utility rather than to establish a clinically optimized staging classifier.

The 14-subcategory representation achieved a pooled ROC-AUC of 0.637, mean cross-validated ROC-AUC of 0.657 +/- 0.061, PR-AUC of 0.634 +/- 0.065, balanced accuracy of 0.624 +/- 0.054, sensitivity for advanced-stage tumors of 0.614 +/- 0.091, and specificity for early-stage tumors of 0.634 +/- 0.076. Its empirical p value was 0.0099 by permutation testing (**Table 2**).

The 14-subcategory representation exceeded the five-system representation (5 features; pooled ROC-AUC 0.623; balanced accuracy 0.585 +/- 0.060) and the random 14-gene-set control (pooled ROC-AUC 0.577; balanced accuracy 0.554 +/- 0.055; empirical p = 0.0297). Higher-dimensional Hallmark 50, KEGG, Reactome, and GO-slim representations achieved pooled ROC-AUC values of 0.667, 0.653, 0.675, and 0.677 using 50, 319, 2052, and 67 features, respectively. These results indicate that the 14-subcategory representation retained progression-related signal beyond random gene-set controls and the coarser five-system representation, although higher-dimensional pathway or ontology baselines achieved somewhat stronger discrimination (**Table 2**).

### 5-D summaries improve compactness and quantitative grounding for LLM-assisted interpretation

To evaluate input suitability for LLM-assisted interpretation, we compared four input modes across 21 colorectal cancer transcriptomic comparisons: organ-level context, gene-level top25, gene-level top100, and 5-D functional summaries. Gene-level top100 required the largest prompt length (6301.2 +/- 189.1 tokens), whereas the 5-D summary required 1309.2 +/- 9.4 prompt tokens, corresponding to a prompt-token ratio of 0.208 +/- 0.006 relative to top100. Gene-level top25 required 2027.3 +/- 51.4 prompt tokens, and organ-level context required 521.1 +/- 7.7 tokens (**Table 3**).

**Table 3.** Automated comparison of four LLM input modes across colorectal cancer transcriptomic comparisons. This table compares organ-level context, gene-level top25, gene-level top100, and 5-D functional-summary inputs across 21 colorectal cancer transcriptomic comparisons. Values are reported as mean +/-SD. Prompt compression versus top100 was calculated as the prompt-token ratio relative to the top100 baseline. Valid numeric anchors denote output values that exactly matched curated numeric evidence provided in the input. Numeric anchor density was calculated as valid numeric anchors per 100 output words. Lower prompt tokens, total tokens, and latency indicate greater efficiency, whereas higher valid numeric anchors and anchor density indicate stronger quantitative grounding.

| Metric | Organ-level context | Gene-level top25 | Gene-level top100 | 5-D summary |
| --- | --- | --- | --- | --- |
| n | 21 | 21 | 21 | 21 |
| Prompt tokens (mean ± SD) | 521.1 ± 7.7 | 2027.3 ± 51.4 | 6301.2 ± 189.1 | 1309.2 ± 9.4 |
| Completion tokens (mean ± SD) | 2247.6 ± 291.3 | 3144.9 ± 614.0 | 3541.8 ± 683.1 | 3357.7 ± 397.3 |
| Total tokens (mean ± SD) | 2768.8 ± 292.9 | 5172.1 ± 611.3 | 9843.0 ± 665.2 | 4666.9 ± 397.6 |
| Latency, s (mean ± SD) | 38.3 ± 8.2 | 42.9 ± 10.4 | 64.7 ± 12.0 | 58.0 ± 18.7 |
| Output words (mean ± SD) | 167.3 ± 19.3 | 257.8 ± 34.9 | 268.9 ± 25.8 | 259.6 ± 43.2 |
| Prompt compression vs top100 (mean ± SD) | 0.083 ± 0.003 | 0.322 ± 0.002 | 1.000 ± 0.000 | 0.208 ± 0.006 |
| Valid numeric anchors (mean ± SD) | 1.81 ± 0.40 | 4.29 ± 2.65 | 3.71 ± 2.92 | 8.48 ± 4.13 |
| Numeric anchor density per 100 words (mean ± SD) | 1.10 ± 0.28 | 1.66 ± 0.98 | 1.39 ± 1.11 | 3.15 ± 1.22 |

The 5-D summary mode also reduced total token use relative to top100 (4666.9 +/- 397.6 versus 9843.0 +/-665.2 total tokens), while producing a completion length comparable to the evidence-bearing gene-level modes. Latency for 5-D summaries (58.0 +/- 18.7 s) was lower than top100 (64.7 +/- 12.0 s) but higher than organ-level context and top25 (38.3 +/- 8.2 s and 42.9 +/- 10.4 s, respectively) (**Table 3**).

Outputs generated from 5-D summaries contained the highest number of valid numeric anchors (8.48 +/-4.13), compared with gene-level top25 (4.29 +/- 2.65), gene-level top100 (3.71 +/- 2.92), and organ-level context (1.81 +/- 0.40). Numeric anchor density was also highest for 5-D summaries (3.15 +/- 1.22 per 100 words), compared with 1.66 +/- 0.98, 1.39 +/- 1.11, and 1.10 +/- 0.28 for top25, top100, and organ-level context, respectively (**Table 3**). These results indicate that the 5-D summary mode was not only more compact than top100 gene-level input, but also more quantitatively grounded in the generated reports.

## Discussion

### Framework-level value: an auditable functional state space

The present framework defines an auditable functional state space between high-dimensional molecular outputs and high-level biological interpretation. Rather than interpreting transcriptomic data only as gene lists, pathway lists, or organ-level descriptions, it reorganizes molecular signals into coordinated activity patterns across five major systems, 14 functional subcategories, and a general infrastructure layer. In this representation, each disease transcriptome or disease-control comparison can be summarized as a functional fingerprint within a shared biological coordinate system. The System 0 infrastructure layer is particularly important because it separates ubiquitous housekeeping machinery from context-dependent disease-state programs. The global distribution, system-level semantic profiles, and supplementary coherence analyses support the view that the framework captures non-random functional neighborhoods while remaining transparent and auditable (**Figures 1-3**, **Table 1, Supplementary Figs. S1-S2**).

### Wound healing illustrates dynamic functional handover rather than a single repair program

The wound-healing analysis shows that physiological tissue repair is not adequately captured by a single monolithic “repair” label. The early multi-system activation peak, delayed A4 activity, secondary C/D/E activity, and phase-specific top-gene turnover indicate that different molecular modules can carry related higher-order functions at different stages of tissue recovery. This distinction between gene-level turnover and system-level functional continuity is a major advantage of the 5-D representation for time-series transcriptomic interpretation. It allows wound healing to be interpreted as a dynamic functional handover among defense, repair, metabolic support, regulatory coordination, and late homeostatic restoration, rather than as a linear activation of one repair pathway (**Figure 4**).

### Sepsis highlights dysregulated host-response organization in blood transcriptomes

The sepsis profile is medically informative because the largest positive shifts were not observed in canonical immune-defense subcategories, but in metabolic-stress and regulatory modules. This pattern should not be interpreted as improved metabolism or more effective regulatory control. In the context of sepsis, increased C3 and D2 more likely reflect stress-adaptive or compensatory overdrive, whereas the reduction of B1-B3 suggests reduced blood-based immune-defense signatures. Because the dataset is based on whole blood, these immune-defense shifts may combine transcriptional changes with leukocyte-composition effects; this limitation should be addressed in future work using deconvolution or single-cell data (**Figure 5**).

### CRC analyses support recurrent tumorization-associated functional fingerprints

The colorectal cancer analysis demonstrates that the framework can recover recurrent functional fingerprints under substantial dataset heterogeneity. Across adenoma, carcinoma/tumor, and metastasis comparisons, repeated elevation of A1 and C2 provides a compact representation of genomic maintenance/repair and biosynthetic/anabolic demand across the CRC spectrum. Persistent A4 positivity and recurrent D2/C3 suppression further suggest that tumor-associated states involve remodeling and damage-management programs together with relative weakening of broader regulatory and metabolic-stress buffering dimensions. These patterns support the use of functional fingerprints as compact disease-state summaries across related pathological stages. Because the included datasets varied in platform, sample source, and disease context, these findings should be interpreted as system-level tendencies rather than as a formal clinical meta-analysis (**Supplementary Table S3, Figure 6**).

### The extended atlas supports scalable cross-disease functional fingerprinting

The extended atlas moves the framework beyond selected case studies toward broader cross-disease comparison. The heterogeneous 14-subcategory patterns across 38 complete case-control datasets suggest that different diseases occupy distinguishable positions in a shared functional state space. Importantly, the atlas should be interpreted as an exploratory scalability test rather than as a harmonized clinical diagnostic atlas. Its main value is to show that diverse disease transcriptomes can be represented as comparable functional fingerprints, providing a basis for future disease-state retrieval, subtype discovery, prognosis modeling, and therapeutic-response prediction (**Figure 7**).

### The TCGA stage benchmark shows information retention under compression, not clinical classifier superiority

The TCGA benchmark provides a controlled test of whether the 14-subcategory representation retains disease-state information in a within-cancer task. The 14-subcategory feature space outperformed the five-system representation and random 14-gene-set controls, supporting the value of biologically organized subcategory resolution. Higher-dimensional pathway and ontology baselines achieved somewhat stronger ROC-AUC values, indicating that the framework should not be interpreted as a superior clinical classifier. Instead, its main value is interpretable information retention under strong dimensional compression. This retention was achieved using only 14 biologically interpretable subcategory features, emphasizing the role of the framework as a compact functional representation rather than a maximal prediction model (**Table 2**).

### LLM input benchmarking shows practical value for human-AI interpretation

The LLM comparison indicates that the five-dimensional summary is not merely a shorter input format. It substantially reduced prompt length relative to top-100 gene-level inputs while increasing valid numeric anchors and numeric anchor density in generated reports. This suggests that structured functional fingerprints can help constrain LLM outputs around quantitative evidence and reduce the risk of diffuse, weakly grounded interpretation. In this sense, the framework provides not only dimensional reduction, but also an interface for human-AI biological interpretation. Nevertheless, this component remains a proof-of-concept and should be validated across additional diseases, prompts, models, expert scoring systems, and retrieval-grounded workflows (**Table 3**).

### Limitations and future directions

Several limitations remain. First, the framework uses a predefined disease-agnostic vocabulary, which improves comparability but may compress disease-specific mechanisms. Second, most analyses are based on bulk transcriptomes, so observed functional shifts may reflect both cellular state changes and cell-composition differences. Third, the wound-healing trajectory was reconstructed from two independent cohorts, so late-stage convergence should be interpreted as relative functional normalization rather than as a fully batch-corrected longitudinal trajectory. Fourth, the sepsis analysis focused on case-control functional reorganization rather than severity, mortality, or treatment-response prediction. Fifth, rule-based assignment improves auditability but requires further stability testing under alternative boundary rules and System 0 filtering strategies. Finally, the LLM analysis evaluates input representation rather than comprehensive model performance. Future work should therefore focus on external validation, cell-type-aware analysis, prognosis and treatment-response prediction, disease-specific refinement, open software implementation, and knowledge-grounded LLM workflows.

Together, these findings support a 5-D functional state space for converting complex disease transcriptomes into compact, auditable functional fingerprints. By preserving interpretable disease-state information while reducing dimensional complexity, the framework provides a basis for cross-disease comparison, systems-level transcriptomic interpretation, and more reliable human-AI analysis of biomedical data.

## Materials and Methods

### Overview of the analytical pipeline and data preprocessing

The analytical workflow centered on the 5-D functional classification framework and included dataset preprocessing, gene-level standardization, probe-to-gene mapping when required, functional gene-set scoring, system-level profiling, and LLM-assisted interpretation (**Figure 1A**). Heterogeneous GEO datasets were converted into unified gene-level expression matrices using rule-based identifier and probe-mapping procedures, with detailed conflict-handling rules provided in **Supplementary Table S4**.

### The 5-D functional state-space framework and classification rules

Biological processes and pathways were assigned to five major functional systems according to dominant physiological objective rather than molecular component, anatomical location, or local pathway membership: Self-Healing and Structural Reconstruction (System A; Repair), Immune Defense (System B; Defense), Energy and Metabolic Homeostasis (System C; Metabolism), Cognitive-Regulatory Control (System D; Control), and Reproduction and Continuity (System E; Reproduction). These systems were subdivided into 14 functional subcategories to preserve essential within-system distinctions while retaining compactness (**Table 1**).

Constitutive molecular machinery required for general cellular viability, such as basal transcription, translation, RNA processing, ribosomal activity, and generic intracellular transport, was assigned to System 0 (Infrastructure). System 0 was used as an infrastructure filter to separate ubiquitous execution machinery from context-dependent physiological objectives. Terms contributing to multiple objectives were assigned using deterministic dominant-objective priority rules (**Figure 1B**).

### Functional scoring and LLM-assisted interpretation

Functional activity was quantified using ssGSEA against the predefined subcategory gene sets derived from the 5-D framework (*21–23*). Subcategory-level enrichment scores were aggregated to parent systems when five-system summaries were required. For LLM-assisted interpretation, the model was provided with structured functional summaries containing system activities, ranked subcategory shifts, and explicit statistical anchors rather than unconstrained gene or pathway lists. Prompt templates and evaluation rules are provided in **Supplementary Table S5**.

### Evaluation design and controlled TCGA benchmark

The framework was evaluated in five linked dimensions: internal coherence of the classification, system-level representation across disease contexts, scalability of cross-dataset disease fingerprinting, controlled retention of progression-related information in TCGA-COAD/READ, and input suitability for LLM-assisted interpretation. Disease applications included wound healing, sepsis, colorectal cancer-related comparisons, and an extended GEO case-control atlas. For the controlled within-disease benchmark, TCGA-COAD/READ tumors were grouped into early-stage (I/II) and advanced-stage (III/IV) disease and represented using 14 subcategories, five parent systems, Hallmark 50, KEGG, Reactome, GO-slim biological processes, and random 14-gene-set controls (*24–30*). Performance was assessed by 5-fold cross-validation using pooled ROC-AUC, fold-level ROC-AUC, PR-AUC, balanced accuracy, sensitivity, specificity, and label-permutation empirical p values.

## Funding

This study was supported by the Fundamental Research Funds for the Central Universities (to J.S.); Sichuan Province Natural Science Foundation Grant 2026NSFSC0606 (to J.S.)

## Author contributions

J.S. conceptualized ideas, designed and managed the project, and wrote and edited the manuscript. F.N. developed the methodology, curated the data, performed the analysis, and wrote and edited the manuscript. J.L. supervised the project and reviewed and edited the manuscript. Y.Z., and K.C. reviewed and edited the manuscript.

## Competing interests

The authors declare that they have no competing interests.

## Data and materials availability

All public transcriptomic datasets analyzed in this study are available from GEO and TCGA, as described in the Materials and Methods and Supplementary Materials. The analysis code is available at https://github.com/exrff/agent-for-human-disease-genes. Processed data tables and intermediate results needed to reproduce the main conclusions are provided in the paper, the Supplementary Materials, and/or the code repository.

## Notes

### Competing Interest Statement

The authors have declared no competing interest.

