## Supplemental file for "A five-dimensional functional state space for fingerprinting disease transcriptomes"

Supplementary Materials
This file includes: Tables S1 to S5；Figs. S1 to S2

**Supplementary Table S1. Phase-specific representative genes supporting functional continuity despite temporal gene turnover during wound healing.**

Representative priority genes were selected from the top activated genes within each functional system at each wound-healing time point. The table summarizes the dominant biological interpretation of each phase, representative priority genes, their main biological meanings, and how temporally changing gene modules support stable higher-order functional objectives. This table supports the interpretation that wound-healing trajectories reflect continuity of physiological objective rather than persistence of identical genes across time.

| Time point / phase | Dominant functional interpretation | Representative priority genes | Main biological meaning | Interpretation for functional continuity |
| --- | --- | --- | --- | --- |
| **Day 1 / Acute priming phase** | Broad inflammatory, vascular, regulatory, and defensive priming precedes overt structural rebuilding. | **NCF1, FGFR1, HEG1, INPPL1, NEUROD1, KALRN, MAP1A** | NCF1 supports phagocyte oxidative burst; FGFR1 and HEG1 suggest reparative/angiogenic and vascular integrity programs; INPPL1 indicates migration-related signaling; NEUROD1/KALRN/MAP1A suggest early neuro-regulatory or signaling participation. | The early peak reflects alarm, defense, vascular adjustment, and regulatory coordination rather than narrow structural repair alone. |
| **Day 3 / Early transition phase** | Gene composition shifts toward inflammation processing, protease control, matrix conditioning, and damage containment. | **CTSC, PI3, AIF1, TIMP1, PDPN, FCER1G, IFI30, BRCA2, UBE2T** | CTSC supports neutrophil protease activation; PI3/elafin indicates anti-protease and anti-inflammatory protection; AIF1 reflects myeloid/endothelial activation; TIMP1 and PDPN indicate matrix and stromal remodeling; BRCA2/UBE2T suggest DNA-damage or proliferative-stress handling. | The molecular module changes from Day 1, but the higher-order objective remains aligned with controlled inflammation, damage handling, and preparation for repair. |
| **Day 7 / Organized repair-competent phase** | Inflammatory execution, epithelial response, matrix control, and immunometabolic support become prominent. | **PI3, CTSC, CD86, C5AR1, CCR1, KRT16, TIMP1, IL6, SLC2A1, HK3, PGAM1** | CD86 indicates antigen-presenting/costimulatory activity; C5AR1/CCR1 support inflammatory recruitment; KRT16 marks wound-edge epithelial stress and re-epithelialization; TIMP1 indicates matrix turnover control; IL6/SLC2A1/HK3/PGAM1 indicate immunometabolic fueling. | Different genes now carry the same broad goals: immune execution, epithelialization, matrix governance, and metabolic support for active repair. |
| **Day 14 / Remodeling phase** | A new remodeling-associated module emerges, centered on stromal activation, extracellular matrix assembly, angiogenic tuning, and morphogenetic programs. | **MDK, CDH11, FKBP10, VASH1, RUNX2, SULF2, BGN, COL6A1, COL6A2, VCAN, ACVR1** | MDK acts as a pan-system regenerative hub; CDH11 suggests fibroblast/myofibroblast and stromal remodeling; FKBP10 and collagen-related genes indicate matrix assembly; VASH1 suggests angiogenic feedback control; RUNX2/VCAN/ACVR1 support remodeling and morphogenetic signaling. | This phase shows near-complete molecular replacement but strong functional coherence with tissue reconstruction and remodeling. |
| **Day 21 / Resolution and homeostatic surveillance phase** | Overt rebuilding declines and is replaced by macrophage-centered cleanup, immune surveillance, iron/nutrient handling, autophagy, and tissue support. | **SIGLEC1, C1QA, CSF1R, SLC40A1, ATG7, IL15RA, RNASE6, STAT3, IGF1, PLTP** | SIGLEC1/C1QA/CSF1R indicate macrophage surveillance and cleanup; SLC40A1 supports iron handling and stromal proliferation; ATG7 indicates autophagy-related homeostatic support; STAT3/IGF1/PLTP suggest late tissue-support and metabolic regulation. | Declining system-level amplitudes do not mean biological inactivity; the wound shifts into cleanup, surveillance, and homeostatic restoration. |
| **Cross-phase conclusion** | System-level trajectories reflect continuity of physiological objective rather than persistence of identical genes. | Gene overlap between adjacent phases is limited, while phase-specific hub genes recur across systems. | Each phase recruits different molecular modules to support defense, inflammatory control, metabolic support, tissue reconstruction, and homeostatic resolution. | The five-dimensional framework captures stable higher-order functions carried by time-varying gene modules. |

**Supplementary Table S2. Full list of colorectal cancer-related disease-versus-normal comparisons.**

This table lists all 21 colorectal cancer-related comparison units included in the broad cross-dataset robustness analysis. Comparisons span adenoma-, carcinoma/tumor-, and metastasis-related contexts. For each comparison, the GEO dataset name, platform, sample source, disease-group size, and control-group size are provided. These datasets support the analysis of recurrent and progression-associated five-dimensional functional patterns across the colorectal cancer spectrum.

| Comparison | Dataset name | Platform | Sample source | Disease group (n) | Control group (n) |
| --- | --- | --- | --- | --- | --- |
| GSE100179 adenoma vs normal | Human Transcriptome Array 2.0 (HTA) from healthy colonic, colorectal adenoma and colorectal cancer tissue | GPL17586 | tissue_biopsy | adenoma (20) | normal (20) |
| GSE198692 adenoma vs normal | Transcriptome of sessile serrated adenoma/polyps shows a relation with MSI-high colorectal cancer and decrease of CDX2 | GPL21185 | tissue_biopsy | adenoma (10) | normal (10) |
| GSE37364 adenoma vs normal | Expression data from human colonic biopsy samples (adenoma-carcinoma) | GPL570 | tissue_biopsy | adenoma (29) | normal (38) |
| GSE41258 adenoma vs normal | Expression data from colorectal cancer patients | GPL96 | tissue_biopsy | adenoma (51) | normal (74) |
| GSE50114 adenoma vs normal | Identification of a gene expression signature in colorectal adenoma in comparison with normal mucosa | GPL6480 | tissue_biopsy | adenoma (37) | normal (9) |
| GSE68468 adenoma vs normal | caArray_notte-00422: Molecular Dissection of Colon Cancer | GPL96 | tissue_biopsy | adenoma (50) | normal (75) |
| GSE77953 adenoma vs normal | Global gene expression analysis of development and progression of colorectal carcinoma | GPL96 | tissue_biopsy | adenoma (17) | normal (13) |
| GSE100179 carcinoma vs normal | Human Transcriptome Array 2.0 (HTA) from healthy colonic, colorectal adenoma and colorectal cancer tissue | GPL17586 | tissue_biopsy | carcinoma (20) | normal (20) |
| GSE125229 carcinoma vs normal | A cancer stem cell hierarchy based on differential ribosomal RNA and protein synthesis capacities [EPHB2] | GPL15207 | tissue_biopsy | carcinoma (24) | normal (4) |
| GSE164191 carcinoma vs normal | Blood based expression of colorectal cancer and healthy controls | GPL570 | peripheral_blood | carcinoma (59) | normal (62) |
| GSE216724 carcinoma vs normal | Colon Cancer Patient-Derived Spheroids and their Stem Cell-Intrinsic mRNA Expression Yield Novel Prognostic Signatures | GPL21185 | tissue_biopsy | carcinoma (58) | normal (6) |
| GSE24514 carcinoma vs normal | Expression data from human MSI colorectal cancer and normal colonic mucosa | GPL96 | tissue_biopsy | carcinoma (34) | normal (15) |
| GSE29638 carcinoma vs normal | Gene level expression profiling of colorectal cancer tissue samples | GPL5175 | tissue_biopsy | carcinoma (48) | normal (2) |
| GSE37364 carcinoma vs normal | Expression data from human colonic biopsy samples (adenoma-carcinoma) | GPL570 | tissue_biopsy | carcinoma (27) | normal (38) |
| GSE68468 carcinoma vs normal | caArray_notte-00422: Molecular Dissection of Colon Cancer | GPL96 | tissue_biopsy | carcinoma (198) | normal (75) |
| GSE70468 carcinoma vs normal | Vitamin D receptor expression and associated gene signature in tumor stromal fibroblasts predict clinical outcome in col | GPL17077 | tissue_biopsy | carcinoma (14) | normal (14) |
| GSE77953 carcinoma vs normal | Global gene expression analysis of development and progression of colorectal carcinoma | GPL96 | tissue_biopsy | carcinoma (17) | normal (13) |
| GSE40367 metastasis vs normal | Gene expression analysis of liver and colon cancer primary tumors and metastasis | GPL570 | tissue_biopsy | metastasis (56) | normal (5) |
| GSE41258 metastasis vs normal | Expression data from colorectal cancer patients | GPL96 | tissue_biopsy | metastasis (67) | normal (74) |
| GSE49355 metastasis vs normal | Specific extracellular matrix remodeling signature of colon hepatic metastases [HG-U133A] | GPL96 | tissue_biopsy | metastasis (19) | normal (18) |
| GSE77953 metastasis vs normal | Global gene expression analysis of development and progression of colorectal carcinoma | GPL96 | tissue_biopsy | metastasis (11) | normal (13) |

**Supplementary Table S3. Summary of colorectal cancer-related comparisons included in the cross-dataset robustness analysis.**

This table summarizes the colorectal cancer-related disease-versus-normal comparisons used for the Figure 6 robustness analysis. Comparisons were grouped into adenoma-, carcinoma/tumor-, and metastasis-related contexts. For each group, the table reports the number of comparison units, unique datasets, disease and control sample sizes, platform coverage, and sample sources.

| Disease group | n comparisons | n unique datasets | Total disease samples | Total control samples | Platforms | Sample sources |
| --- | --- | --- | --- | --- | --- | --- |
| Adenoma | 7 | 7 | 214 | 239 | GPL17586; GPL21185; GPL570; GPL96; GPL6480 | tissue_biopsy |
| Carcinoma | 10 | 10 | 499 | 249 | GPL17586; GPL15207; GPL570; GPL21185; GPL96; GPL5175; GPL17077 | tissue_biopsy; peripheral_blood |
| Metastasis | 4 | 4 | 153 | 110 | GPL570; GPL96 | tissue_biopsy |

**Supplementary Table S4. GEO preprocessing and probe-to-gene mapping rules.**

This table summarizes the rule-based preprocessing and probe-to-gene mapping workflow used to convert heterogeneous GEO expression matrices into standardized gene-level expression matrices. Each row describes a specific input condition or conflict type, the corresponding operational rule, the conflict-resolution policy, the rationale for reproducibility, and the relevant code basis. The rules were designed to ensure deterministic handling of malformed matrices, unsupported study types, probe-like identifiers, platform annotation conflicts, ambiguous gene-symbol fields, many-to-one probe mappings, and post-mapping sanity checks before downstream ssGSEA scoring.

| Stage | Condition or conflict type | Operational rule | Conflict-resolution policy | Rationale for reproducibility | Code basis |
| --- | --- | --- | --- | --- | --- |
| Series-matrix validation | Malformed or non-usable GEO matrix | Require a `!series_matrix_table_begin`/`!series_matrix_table_end` block with at least one expression-data row beyond the header. | Stop preprocessing if the table block is missing or header-only. | Prevents downstream parsing of empty, malformed, or metadata-only GEO records. | src/agent/geo_parsing.py:8-101 |
| Series-matrix validation | Unsupported study type or organism | Reject SuperSeries without a usable matrix, non-expression studies (for example ChIP-seq, ATAC-seq, methylation/CNV), mouse datasets, and datasets with fewer than 6 samples. | Stop preprocessing and record the exclusion reason. | Restricts the pipeline to analyzable human expression datasets with minimal comparison power. | src/agent/geo_parsing.py:13-26, 71-97 |
| Series matrix parsing | Quoted text, mixed cell types, or empty rows | Use the first table column as the row index (`probe_id`), coerce all sample columns to numeric values, and drop rows that are all-NaN after coercion. | Retain partially valid rows but remove rows with no usable numeric signal. | Standardizes heterogeneous GEO matrix encodings into a numeric expression matrix. | src/agent/geo_parsing.py:135-166 |
| Identifier typing | Row IDs may already be genes or may still be probes | Infer `probe`, `gene`, or `unknown` from the first 300 row IDs using regex heuristics. A probe-like ratio >=0.10 is classified as `probe`; strong gene-like evidence (>=0.6) with probe-like ratio <0.10 is classified as `gene`; an intermediate probe-like ratio >=0.05 is also treated as `probe`. | Use rule-based typing before deciding whether GPL annotation is required. | Avoids unnecessary remapping of already gene-level matrices while preserving safety for probe-like inputs. | src/agent/geo_parsing.py:169-213 |
| Safety override | Matrix looks gene-like overall but still contains obvious probe signatures | If the matrix was inferred as `gene` but the empirical probe-like ID ratio is >=0.08, force probe-to-gene mapping when GPL annotation is available; otherwise abort preprocessing. | Prefer a conservative forced remapping rather than trusting a potentially misclassified matrix. | Reduces false pass-through of probe matrices that superficially resemble gene symbols. | src/agent/analysis_nodes_data.py:255-279 |
| Direct gene-level pass-through | Input matrix is already gene-level | Skip GPL mapping, strip blank gene IDs, and collapse duplicate gene IDs by arithmetic mean. | Many-to-one duplicates at the gene level are averaged into a single gene row. | Ensures a unique gene index before downstream ssGSEA scoring. | src/agent/analysis_nodes_data.py:281-286 |
| GPL discovery | Platform annotation file must be matched to the GEO series | Read `!Series_platform_id` from the series matrix, search `data/gpl_platforms/` first, then the dataset folder, and prefer files whose names start with the detected GPL accession. | If no GPL file is found for a probe-like matrix, stop preprocessing. | Pins probe mapping to the intended platform rather than using a generic annotation table. | src/agent/geo_parsing.py:104-132; src/agent/analysis_nodes_data.py:287-291 |
| GPL table parsing | GPL files vary between family.soft tables and plain tabular annotations | Parse from `!platform_table_begin` when present; otherwise fall back to the first non-comment, non-control line. | Raise an error if no tabular annotation block can be located. | Supports multiple official GPL file layouts without manual editing. | src/agent/geo_parsing.py:216-259 |
| Probe-column selection | Probe identifiers may be stored under different header names | Prefer columns named `ID`, `PROBE_ID`, or `PROBEID`; otherwise use the first GPL table column as the probe identifier. | Normalize the selected column to a standard `probe_id` field. | Provides a deterministic rule for heterogeneous vendor-specific GPL headers. | src/agent/geo_parsing.py:261-264 |
| Preferred gene-symbol column selection | GPL annotation may provide several candidate gene-symbol columns | Prioritize explicit gene/symbol headers (`Gene Symbol`, `SYMBOL`, `gene_assignment`, etc.); if none exist, fall back to the first column containing the token `gene`. | Normalize the chosen column to a standard `gene_symbol` field. | Avoids ad hoc manual column picking across platforms. | src/agent/geo_parsing.py:266-281, 332-335 |
| Special-case override for `SPOT_ID` | One-to-one column conflict: `ID` stores Ensembl IDs while `SPOT_ID` stores the human-readable gene symbol | When `ID` is predominantly Ensembl-like, `SPOT_ID` is predominantly symbol-like, and the initially selected gene column is absent or still Ensembl-like, replace the gene-symbol source with `SPOT_ID`. | Override the earlier gene-column choice in favor of `SPOT_ID`. | Handles GPL families where the biologically interpretable symbol is not in the nominal gene column. | src/agent/geo_parsing.py:283-305 |
| NanoString-like fallback | Explicit gene column missing, but `SPOT_ID` itself behaves like a gene-symbol column | If `SPOT_ID` is symbol-like, mostly non-numeric, sufficiently unique (>0.15 unique ratio), and not dominated by low-information categorical labels, treat `SPOT_ID` as the gene-symbol source. | Accept `SPOT_ID` as the best available mapping column. | Rescues platforms that encode symbols directly in `SPOT_ID` without a conventional `Gene Symbol` header. | src/agent/geo_parsing.py:306-330 |
| Free-text annotation fallback | No explicit gene-symbol column is available | Select the longest annotation-like column among `SPOT_ID*`, `description`, or `transcript` fields, then extract the best candidate symbol from free text using curated regex patterns or a plain-symbol fallback. | Derive one best-effort symbol per probe from annotation text; fail only if no annotation-like column exists. | Recovers usable probe annotations from verbose GPL descriptions rather than discarding the platform. | src/agent/geo_parsing.py:336-355, 376-393 |
| One-to-many conflict handling | A single probe annotation cell lists multiple possible genes | Keep only the primary mapping block before `///`, split `accession // SYMBOL // description` patterns, and test candidates in priority order `[1, 0, 2]`. Reject source-like tokens (`ENSEMBL`, `REFSEQ`, `HGNC`, etc.) and accession-like tokens before accepting a symbol-like token. | Collapse each ambiguous probe record to one best symbol; if no plausible symbol survives, drop the record. | Prevents uncontrolled fan-out from one probe to multiple genes while keeping a deterministic primary mapping rule. | src/agent/geo_parsing.py:396-435 |
| Invalid-symbol filtering | Placeholder or low-information annotation values | Drop null/empty symbols and filter low-information labels such as `---`, `null`, `nan`, `coding`, `multiple_complex`, `multiple`, `complex`, `control`, and `main`. | Remove ambiguous rows before matrix aggregation. | Improves symbol specificity and prevents non-biological labels from entering the gene matrix. | src/agent/geo_parsing.py:357-373 |
| Many-to-one conflict handling | Multiple probes map to the same gene | Inner-join the probe expression matrix with the cleaned probe-to-gene mapping, drop `probe_id`, then aggregate all probes assigned to the same `gene_symbol` by arithmetic mean. | Represent each gene by a single averaged expression profile. | Produces a unique gene-level matrix required by the downstream ssGSEA workflow. | src/agent/geo_parsing.py:438-442 |
| Post-mapping sanity check | Suspiciously small mapped gene matrix after probe-like input | If the mapped gene matrix contains fewer than 200 genes while the probe-like ratio is >=0.08, flag the result as likely platform/annotation mismatch and stop preprocessing. | Abort rather than continue with a likely incorrect mapping. | Prevents silently propagating catastrophic GPL mismatches into downstream biological interpretation. | src/agent/analysis_nodes_data.py:293-307 |

**Supplementary Table S5. LLM prompt design, control mechanisms, and evaluation metrics.**

This table summarizes the prompt architecture, embedded control instructions, implementation notes, and evaluation metrics used in the LLM-assisted interpretation workflow. It includes the main disease-analysis prompt scaffold, five-dimensional classification principles, mode-focus JSON structure, additional interpretation constraints, API-level implementation details, four-arm CRC evaluation prompts, automated efficiency metrics, evidence-grounding metrics, format-compliance checks, and optional blinded-review scaffolds. The table documents how LLM outputs were constrained, grounded in quantitative evidence, and evaluated for compactness, numerical anchoring, and reproducibility.

| Category | Item | What is provided to the LLM | Anti-hallucination or control mechanism | Evaluation or operational note | Code basis |
| --- | --- | --- | --- | --- | --- |
| Prompt template | Analysis-strategy router | Dataset metadata (`dataset_id`, English name, Chinese name, disease type, description) plus a closed set of candidate strategies: `case_control`, `subtype_comparison`, `time_series`, and `correlation`. | Restricts the model to a closed decision set and requires JSON-only output with predefined keys (`strategy`, `reasoning`, `confidence`, `secondary_analyses`, `key_focus`). | Used for task routing rather than narrative generation. | src/agent/prompts.py:36-47; data/prompts/analysis_strategy.txt |
| Prompt template | Visualization-strategy planner | The selected analysis strategy plus structured data characteristics, together with a whitelist of admissible chart types. | Forces the response into JSON and a fixed visualization vocabulary; explicitly asks for downgrade reasons when metadata are incomplete. | Used to recommend figure panels while preserving mode-specific compatibility. | src/agent/prompts.py:50-65; data/prompts/visualization_strategy.txt |
| Prompt template (main pipeline) | Main result-interpretation scaffold | Dataset metadata, the five-dimensional system schema (Systems A-E and subcategories), a compact score summary, and full structured statistical results injected into the prompt body. | Grounds the model in explicit quantitative inputs rather than free-form disease description alone. | This is the core narrative-generation prompt used by the main disease-analysis pipeline. | src/agent/prompts.py:74-99; data/prompts/result_interpretation.txt |
| Prompt-embedded system instruction (main pipeline) | Five-dimensional short principles | An appended rule block that states the objective-driven classification principles, including System A-E priority rules and the explicit assignment of constitutive shared machinery to System 0. | Reduces category drift by making the intended ontological boundaries explicit before interpretation. | Loaded from a separate text file and appended automatically by `_load_compact_principles()`. | src/agent/prompts.py:32-35, 129; data/prompts/System_Classification_Principles.txt |
| Prompt-embedded system instruction (main pipeline) | Mode Focus JSON | A structured JSON block summarizing the current analysis mode, activated/suppressed systems, group counts, top up/down subcategories, and strongest positive/negative associations. | Forces attention onto mode-relevant evidence and discourages generic disease summaries that ignore the actual statistical design. | Constructed dynamically from the statistical results dictionary. | src/agent/prompts.py:68-72, 120-128 |
| Prompt-embedded system instruction (main pipeline) | Extra interpretation requirements | A fixed instruction block requiring discussion of both up- and down-regulated systems/subcategories, explicit between-group comparison when groups exist, evidence-vs-hypothesis separation, and 3-5 testable conclusions. | Separates observed evidence from speculation and forces the model to stay tied to the measured comparison structure. | Appended after the main interpretation prompt and Mode Focus JSON. | src/agent/prompts.py:100-119 |
| Implementation note (main pipeline) | API-level packaging of instructions in the production wrapper | The production client sends a single `user` message containing the prompt body; there is currently no separate API-level `system` role in the wrapper. | The effective 'system instructions' are implemented as prompt-embedded rule blocks rather than a standalone system message. | Important for exact reproducibility of the main disease-analysis reports. | src/agent/llm_client.py:105-123 |
| Prompt template (Experiment 3.5) | Shared system prompt for CRC batch evaluation | A fixed expert-role instruction requiring evidence-grounded biological interpretation, explicit distinction between observed evidence and inferred mechanisms, limitation statements when evidence is insufficient, terminology neutrality, and concise writing. | Explicitly forbids unsupported gene mutations, clinical outcomes, drug-response claims, and internal framework jargon in the final report. | Used identically across all four CRC input modes to isolate the effect of input representation rather than prompt persona. | scripts/run_experiment34_crc_batch_llm_evaluation.py (SYSTEM_PROMPT); results/experiment34_llm_crc_batch_en/prompt_protocol.md |
| Prompt template (Experiment 3.5) | Shared output-format constraint | A fixed three-section Markdown output structure: `Core Findings`, `Systemic Interpretation`, and `Conclusion`. | Requires 2-3 evidence-bearing bullets in the findings section, explicit acknowledgement of weak or mixed support, and cautious downstream inference. | Used to standardize downstream comparison across 21 CRC transcriptomic comparisons and four input modes. | scripts/run_experiment34_crc_batch_llm_evaluation.py (OUTPUT_FORMAT_BLOCK); results/experiment34_llm_crc_batch_en/prompt_protocol.md |
| Input mode (Experiment 3.5) | Organ-level context arm | Only experimental-design context: dataset ID, dataset name, comparison type, disease context, platform, sample source, and case/control sample sizes. | The task explicitly instructs the model to provide only cautious high-level expectations and to state when no quantitative molecular evidence is available. | Serves as the minimal-context baseline for the CRC batch evaluation. | scripts/run_experiment34_crc_batch_llm_evaluation.py (_build_user_prompts); results/experiment34_llm_crc_batch_en/prompt_protocol.md |
| Input mode (Experiment 3.5) | Gene-level top25 arm | Top 25 up-regulated and top 25 down-regulated genes, each annotated with Cohen's d, mean difference, and q value, together with the same dataset brief used in the organ-level arm. | The task instructs the model to prioritize the strongest quantitative patterns and to separate direct observations from downstream inference. | Functions as the compact gene-list baseline in the 4-arm CRC comparison. | scripts/run_experiment34_crc_batch_llm_evaluation.py (_build_user_prompts, _save_prepared_artifacts) |
| Input mode (Experiment 3.5) | Gene-level top100 arm | Top 100 up-regulated and top 100 down-regulated genes, each annotated with Cohen's d, mean difference, and q value, together with the same dataset brief used in the organ-level arm. | The task preserves the same evidence-grounding instructions as the top25 arm, but tests performance under a larger context window. | Functions as the extended dense gene-list baseline in the 4-arm CRC comparison. | scripts/run_experiment34_crc_batch_llm_evaluation.py (_build_user_prompts, _save_prepared_artifacts) |
| Input mode (Experiment 3.5) | Five-dimensional summary arm | A structured system-level and subcategory-level summary containing biologically translated functional-program names plus Cohen's d, mean difference, and q value for each shift. | The prompt presents explicit quantitative anchors while requiring the model to avoid internal framework codes in the final report. | Tests whether a structured functional middle layer improves compactness and quantitative grounding relative to gene-list baselines. | scripts/run_experiment34_crc_batch_llm_evaluation.py (_format_system_subcat_table, _build_user_prompts) |
| Automated metrics (Experiment 3.5) | Efficiency metrics | Post hoc metrics computed after generation: `prompt_tokens`, `completion_tokens`, `total_tokens`, `latency_seconds`, and `prompt_token_compression_rate_vs_gene100`. | Not a prompt constraint by itself, but quantifies the cost of each input representation under a fixed interpretation task. | Used in Table 4 to compare four input modes across 21 CRC transcriptomic comparisons. | scripts/run_experiment34_crc_batch_llm_evaluation.py (_calc_automated_metrics); results/experiment34_llm_crc_batch_en/automated_metrics.csv |
| Automated metrics (Experiment 3.5) | Evidence-grounding metrics | Post hoc metrics: `valid_numeric_anchor_hits`, `numeric_anchor_total`, `numeric_anchor_coverage`, and `numeric_anchoring_density_per_100_words`. | Quantifies whether the generated report reuses explicit numerical evidence from the prompt rather than drifting into unsupported generalities. | Anchor sets differ by arm: sample-count anchors for organ-level input, quantitative gene-level anchors for top25 and top100, and system/subcategory effect-size anchors for the five-dimensional summary. | scripts/run_experiment34_crc_batch_llm_evaluation.py (_build_numeric_anchor_map, _calc_automated_metrics); results/experiment34_llm_crc_batch_en/automated_metrics.csv |
| Automated metrics (Experiment 3.5) | Format-compliance metric | A binary check `has_required_sections` verifies whether the returned report contains the exact three mandated Markdown section headers. | Distinguishes factual drift from prompt-following failure and ensures that all arms are compared under the same output structure. | All four modes reached 100% required-section compliance in the batch CRC evaluation. | scripts/run_experiment34_crc_batch_llm_evaluation.py (_has_required_sections); results/experiment34_llm_crc_batch_en/automated_metrics.csv |
| Human-review rubric (Experiment 3.5) | Blinded domain-review dimensions | Not an input to generation; instead, blinded review packets are scored on `Pathological Alignment`, `Interpretive Coherence`, `Evidentiary Discipline`, and `Readability & Utility`. | Keeps biological plausibility assessment separate from automated token and numeric-anchor metrics, and does not require reviewers to endorse the upstream five-dimensional framework itself. | Prepared as a blinded review scaffold for optional human evaluation of the CRC batch outputs. | results/experiment34_llm_crc_batch_en/evaluation_rubric.md; results/experiment34_llm_crc_batch_en/expert_blinded_review_instructions_zh.md |
| Operational safeguard | Trace capture, fallback handling, and blinded packet generation | The evaluation pipeline records prompt text, response text, attempted models, token usage, latency, response paths, and blinded review packets with separate mapping keys. | Provides an audit trail for unsupported outputs, preserves fallback transparency, and separates raw response identity from blinded review order. | Part of the reproducibility and QA framework around the 4-arm CRC batch evaluation rather than part of the prompt itself. | scripts/run_experiment34_crc_batch_llm_evaluation.py; results/experiment34_llm_crc_batch_en/llm_calls.csv; results/experiment34_llm_crc_batch_en/expert_review_blinding_key.csv |

**Supplementary Figures**

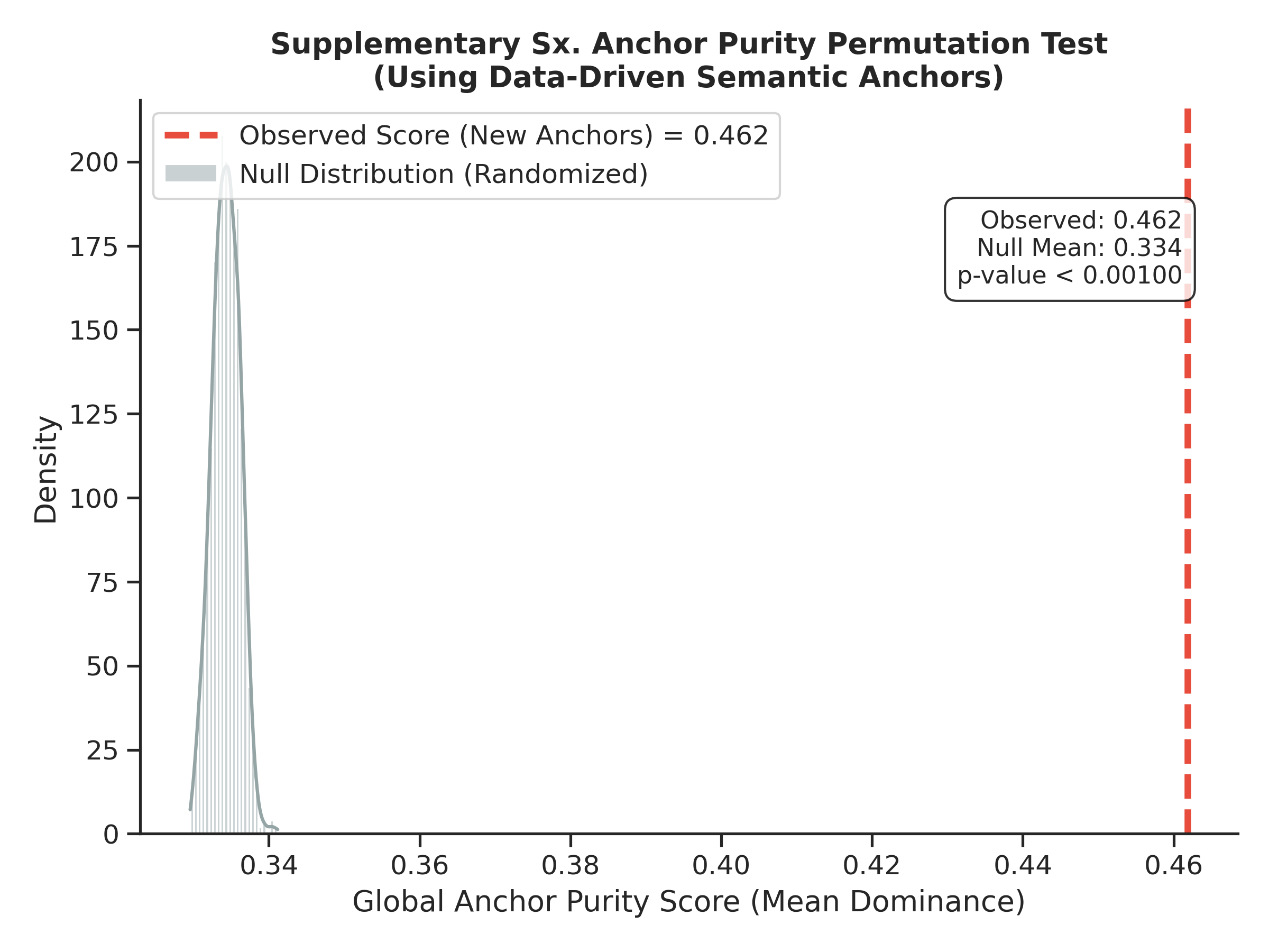

**Supplementary Fig. S1. Anchor purity permutation test for semantic coherence of system assignments.**

For each ontology term, the combined “Name + Definition” text was scanned for matches to predefined semantic anchor keyword sets. Anchor hits were converted into binary presence values per semantic domain. For each functional system, anchor hit counts were normalized across anchor domains, and system purity was defined as the maximum normalized anchor proportion. The global purity score was calculated as the mean purity across systems. The null distribution was generated by shuffling system labels 1,000 times while preserving system sizes. The observed global purity lies in the extreme tail of the null distribution, indicating that system assignments exhibit semantic concentration beyond random partitioning.

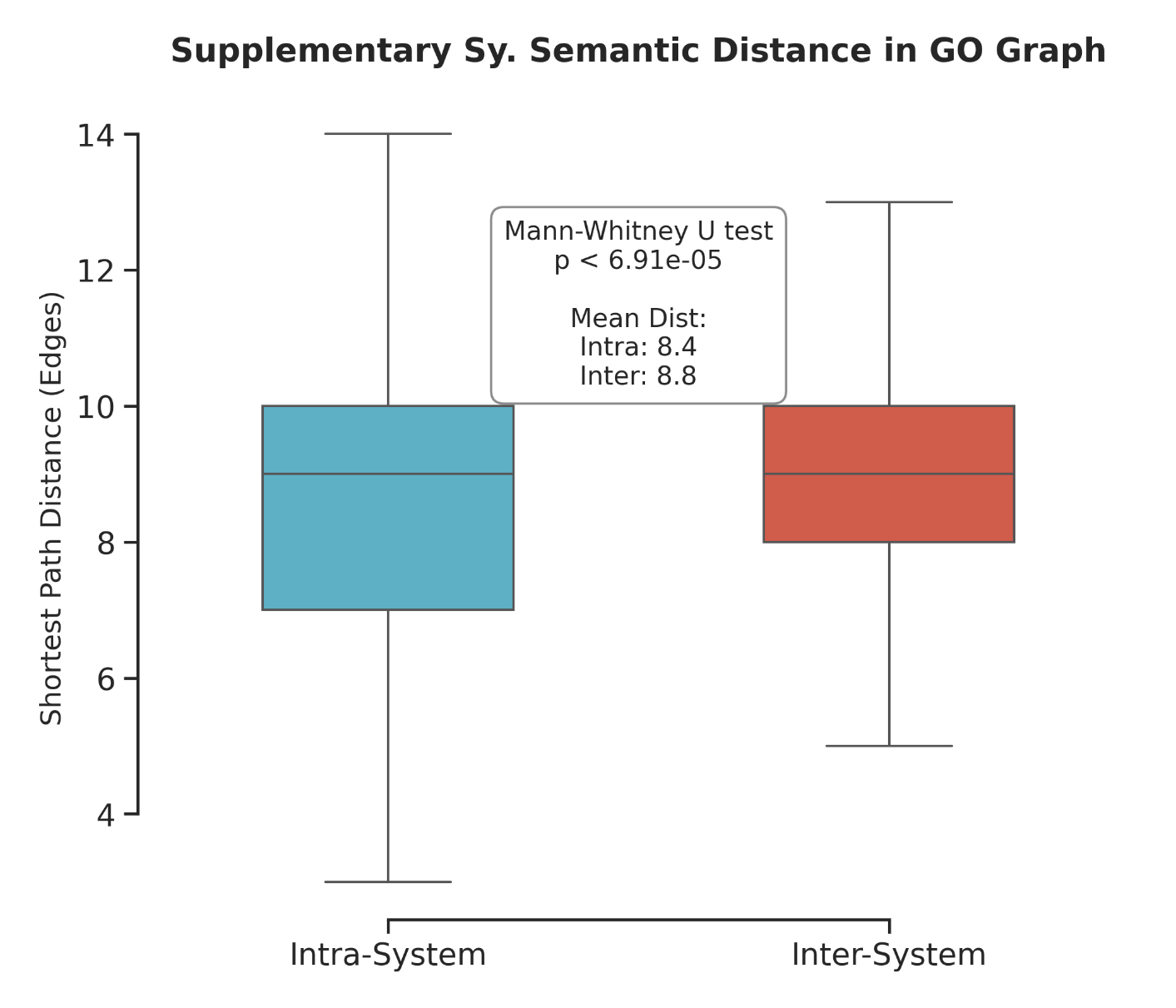

**Supplementary Fig. S2. GO graph semantic distance within versus between functional systems.**

The GO Biological Process graph was constructed from go-basic.obo using is_a edges only and treated as an undirected graph. For each pair of terms, semantic distance was calculated as the shortest path length in the graph. Intra-system term pairs showed significantly shorter graph distances than inter-system term pairs, supporting that the five functional systems correspond to coherent neighborhoods in the GO topology. Unreachable term pairs were excluded from the analysis.
